# Single CAR-Dual target: Intracranial Delivery of Anti-PD-L1 CAR T Cells Effectively Eradicates Glioma and Immunosuppressive Cells in the Tumor Microenvironment

**DOI:** 10.1101/2024.11.22.624897

**Authors:** Maria J. Ulloa-Navas, Yan Luo, Jesus E Sanchez-Garavito, Yaqing Qie, Vanessa K Jones, Mieu M Brooks, Manuela Aramburu-Berckemeyer, Tanya Hundal, Martha E. Gadd, Cristina Gomez-Palmero, Andrew D. Brim, Rocio Rivera-Valentin, Aleeshba N. Basil, Frederick Q. Tutor-New, Jacob B. Hirdler, Sarosh R Irani, Roxana Dronca, Mohamed A Kharfan-Dabaja, Yingxue Ren, Haidong Dong, Hong Qin, Alfredo Quinones-Hinojosa

## Abstract

With the goal to overcome the limited treatment options and poor prognosis of glioblastoma (GBM), we have developed a PD-L1-targeting CAR T cell therapy, MC9999. In vitro experiments with MC9999 CAR T cells derived from GBM patients exhibited potent, antigen-specific cytotoxicity against autologous tumor cells and immunosuppressive cells within the tumor microenvironment (TME). In an orthotopic GBM model using patient-derived brain tumor-initiating cells, the intracranial delivery of MC9999 CAR T cells eradicated established tumors and improved survival. Single-cell RNA sequencing indicated that MC9999 CAR T cells activate interferon pathways, leading to GBM cell apoptosis. Multi-immunohistochemistry confirmed localized PD-L1 expression on tumor cells and TME-residing macrophages, but not in neurons or glia in patient tissue. The local delivery of MC9999 CAR T cells may be a safe, effective approach for simultaneously targeting PD-L1-positive GBM and its immunosuppressive TME and a strategy to overcome immune evasion and enhance the therapeutic potency of CAR T cell therapy against GBM.

## INTRODUCTION

Adult diffuse gliomas are a heterogeneous family of primary tumors that arise within the central nervous system^<u>1</u>^. Among these, the most aggressive are high-grade gliomas (HGG), which include oligodendrogliomas (IDH mutant and 1p/19q co-deleted, WHO grade 3 or anaplastic), astrocytomas (IDH mutant, WHO grade 3 or anaplastic, and WHO grade 4), and glioblastomas (IDH wild type, WHO grade 4)^<u>2</u>^. Grade 4 tumors, such as glioblastoma (GBM) and grade 4 astrocytoma (AS), are common and aggressive primary brain cancers in humans. Despite the standard treatment regimen—maximal surgical resection followed by chemoradiation—median survival for GBM patients remains dismally approximately 14.6 months^<u>3,4</u>^.

The limited success of current treatments can be attributed to several key challenges, including tumor cell heterogeneity and the highly immunosuppressive tumor microenvironment (TME). Tumor heterogeneity in GBM is well-documented, with different subpopulations of cells that shift phenotypically between glial progenitor, neuronal progenitor, and mesenchymal-like states depending on microenvironmental cues^<u>5–9</u>^. These subpopulations express distinct molecular markers such as PDGFRA (platelet derived growth factor receptor A), IDH1/2 (Isocitrate dehydrogenase 1/2), EGFR (epidermal growth factor receptor), and NF1 (neurofibromin), which complicate efforts to target the entire tumor with a single therapeutic agent^<u>10,11</u>^.

Programmed Cell Death ligand 1(PD-L1/CD274/B7-H1), an immune checkpoint molecule, is often hijacked by both tumor cells and immunosuppressive elements within the TME to evade immune destruction. By engaging with PD-1 on T cells, PD-L1 promotes T cell exhaustion, significantly weakening the anti-tumor response ^<u>12</u>^. Based on this observation, in gliomas, particularly GBM and AS, PD-L1 is expressed on both tumor associate macrophage(TAMs) and tumor cells, where it inhibits T cell proliferation through the dephosphorylation of Zap70, a key signaling molecule in T cell activation^<u>13</u>^. Monoclonal antibodies (mAbs) targeting PD-L1 have shown limited efficacy in GBM^<u>14,15</u>^, likely as they can block binding without providing killing capability against antigen-expressing tumor cells. By contrast, CAR T cells can directly kill the cell expressing the antigens, while retaining antigen specificity of mAbs.

In addition, the GBM TME plays a pivotal role in promoting immune evasion. The TME is primarily composed of TAMs and microglia, which exhibit a dual role by contributing to immunosuppression and promoting tumor growth and invasion^<u>16</u>^. TAMs secrete immunosuppressive cytokines such as IL-10 and express immune checkpoint molecules like PD-L1, which induces T cell exhaustion and inhibits anti-tumor immune responses. Furthermore, TAMs express molecules such as MARCO (Macrophage receptor with collagenous structure), HMOX1 (Heme oxygenase 1), ApoE (Apolipoprotein E), NLRP1 (NLR family pyrin domain containing 1), and CD73 (NT5E-5’-nucleotidase ecto), which support tumor progression, invasion, and stemness, while also enhancing immune suppression^<u>17–19</u>^.

Chimeric antigen receptor (CAR) T cell therapy has emerged as a novel cancer immunotherapy, revolutionizing treatment for various types of B-cell non-Hodgkin lymphoma, acute lymphoblastic leukemia, chronic lymphocytic leukemia, and multiple myeloma^<u>20–27</u>^. However, CAR T cell efficacy in GBM is limited due to tumor heterogeneity^<u>28–30</u>^, which requires CAR designed to target multiple antigens. Additionally, the immunosuppressive TMEs in GBM often restricts sustained CAR T cell activity, leading to only transient tumor reduction and impeding the recruitment of additional endogenous active T cells to the tumor site.

Here, we leverage our novel PD-L1-trageting CAR construct (MC9999 CAR) which exhibits potent efficacy by targeting both the tumor cells and the immunosuppressive cell in the TME^<u>31</u>^. This CAR is designed to achieve precise antigen specific cytotoxicity gainst GBM in patient-derived models using intracranial CAR T cell delivery for GBM, and it has demonstrated robust anti-tumor activity in both in vitro assays and in vivo models using GBM patient-derived primary tumor cells. Notably, single-cell RNA sequencing (scRNA-seq) analysis reveals that upon tumor engagement, both CD4+ and CD8+ CAR T cells activate interferon signaling pathways upon tumor engagement, suggesting a strong and coordinated immune response. In addition, GBM patient-derived MC9999 CAR T cells efficiently target autologous immunosuppressive macrophages from the TME, dismantling the tumor’s protective immunosuppressive shield, facilitating the recruitment and activation of endogenous T cells. This dual action—targeting tumor cells and reprogramming the TME—paves the way for a more sustained and comprehensive anti-tumor response. Finally, we demonstrate safety for intracranial delivery of this product for human translation by showing that PD-L1 is only expressed in the target cells and not in neurons, glia or vessels.

## RESULTS

### Anti-PD-L1 (MC9999 CAR-T) cells display high antigen specificity against PD-L1 expressing GBM tumor *in vitro* and *in vivo*

To validate MC9999 CAR T-cells specific cytotoxicity towards PD-L1-expressing targets in a GBM model, we co-cultured healthy donor-derived MC9999 CAR T cells or non-CAR T cells withthe human glioma cell line LN229. This cell line was genetically modified to either overexpress PD-L1 (LN229 PD-L1 OE, Supplementary Fig. 1A) or knock out PD-L1 (LN229 PD-L1 KO).

CD107a expression, a marker of immune cell activation and cytotoxic degranulation, was increased 20-fold (from 1.43% in Non-CAR T to 28.1% in MC9999 CAR T cells) in response to LN229 PD-L1 OE, but not in PD-L1 KO or control groups **(Figure 1A)**. T cell cytotoxicity was confirmed by measuring granzyme B release, which was significantly higher in the MC9999 CAR T treated with LN229 PD-L1 OE cells (N=3, P<0.0001) **(Figure 1B)**. Additionally, a cellular impedance-based cytotoxicity assay demonstrated direct cytotoxic effects of MC9999 CAR T cells on target cell populations, with a sharp reduction in the cell index (inversely related to impedance) observed exclusively in the LN229 PD-L1 OE cells upon addition of MC9999 CAR T cells **(Figure 1C)**.

**Figure 1.**
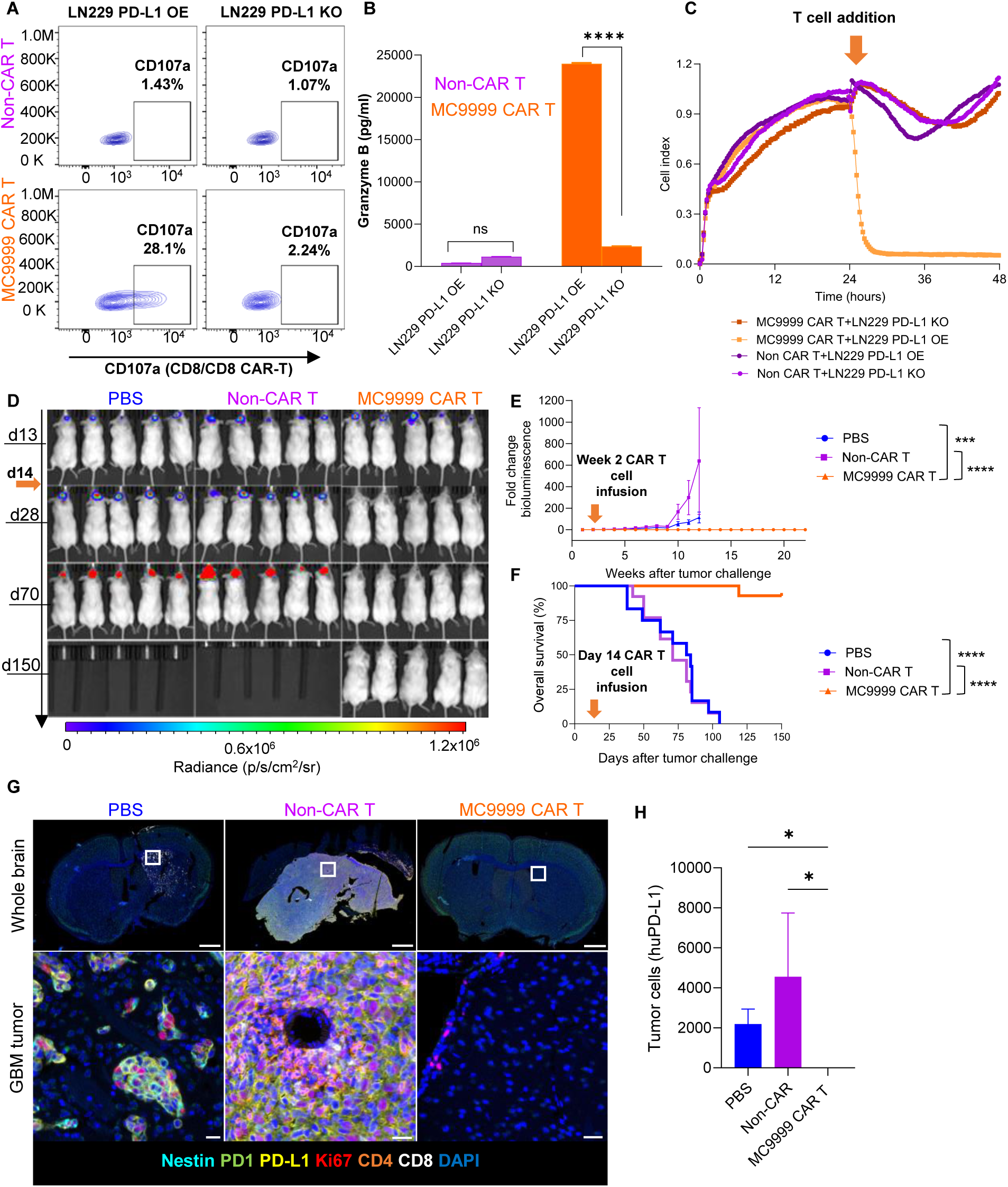
MC9999 CAR T cells display high specificity for PD-L1 expressing cells. **A** LN229 PD-L1-OE and KO cell lines were incubated with MC9999 CAR T cells and Non-CAR T cells from the same healthy donor. D The degranulation assay shows CD107a is highly expressed on CD8+ MC9999 CAR T cells when incubate with LN229 PD-L1 OE cells. **B.** Granzyme B granule secretion is significantly increased when MC9999 CAR T cells were co-cultured with GBM cells expressing the target antigen (n=3; P<0.0001). **C**. After T cell addition (orange arrow), only LN229 PD-L1-OE cells show a decrease in cell index, demonstrating the specific effect of MC9999 CAR T cells on target cells expressing PD-L1 (n=3 per condition). **D.** Bioluminescence-based imaging of tumor cell specificity showed that tumors were depleted without recurrence in the MC9999 CAR T-treated group after 150 days, whereas tumor developed in PBS or Non-CAR T groups. **E.** Decreased bioluminescence and **F.** increased overall survival were observed in MC9999 CAR T-treated groups, but not in Non-CAR T or PBS control groups (n=16 per group, P<0.001). **G.** multi-immunohistochemistry (mIHC) analysis of residual antigen expression in tissue showed proliferative tumor cells expressing PD-L1 and nestin in control groups but not in CAR T treatment group. **H.** PD-L1 expression showing % of remaining tumor cells was significantly lower in CAR T treatment group at endpoint (n=4 per group, P<0.05). Scale bars: 1 mm, magnification 50 µm.

Next, we evaluated the efficacy of MC9999 CAR T cells in vivo using an orthotopic GBM xenograft model whereLN229 PD-L1 OE cells were further transduced with a luciferase reporter to facilitate tumor tracking. Two weeks after tumor implantation and post-confirmation of tumor burden in all animals, a single intratumoral injection of MC9999 CAR T cells, but not non-CAR T cells from the same donor or PBS as a control, led to tumor eradication within two weeks. Further, no tumor recurrence was detected during a 150-day follow-up period. In contrast, the continued tumor growth in both control groups (N=16 per group, P<0.0001) **(Figure 1D, E)** resulted in significantly extended survival for MC9999 CAR T-treated mice **(Figure 1F)**. Consistent with these findings, multi-immunohistochemical (m-IHC) analysis revealed a significant reduction in both human PD-L1-positive tumor cells and Nestin positive cells tumor steam cells in the MC9999 CAR T treated group compared to both control groups (N=5 per group, P<0.05) **(Figure 1G, H; Figure S2)**. Importantly, we also stained for human Nestin, a tumor marker that could indicate antigen (PD-L1) loss. No residual tumor cells were detected in the MC9999 CAR T-treated animals, suggesting complete tumor eradication without antigen escape. These findings provide preclinical evidence that MC9999 CAR T cells effectively and specifically target PD-L1-expressing GBM.

### Anti-PD-L1 (MC9999 CAR-T) cells exhibit high anti-tumor efficacy in GBM patient derived tumor cells in vitro and in vivo

Next, we evaluated the anti-tumor efficacy of MC9999 CAR T cells in patient-derived primary GBM models using cells (BTICs) directly from a GBM (QNS108) patient **(Figure 2A; Table S2).** Given that PD-L1 is a dynamic target in glioma^<u>32–34</u>^, and its expression can decrease under non-stimulating culture conditions (e.g., absence of immune cells or external stimuli like cytokines, radiation, or hypoxia)^<u>35–40</u>^, we transduced the QNS108 BTIC line to stably overexpress PD-L1 **(∼98.9%, Figure S1A)**. Subsequently we further transduced the cell line to express luciferase and GFP (QNS108-PD-L1-GFP-Luc **(Figure 2B)**. This engineered cell line was used as a target to model GBM for in vitro and in vivo studies.

**Figure 2.**
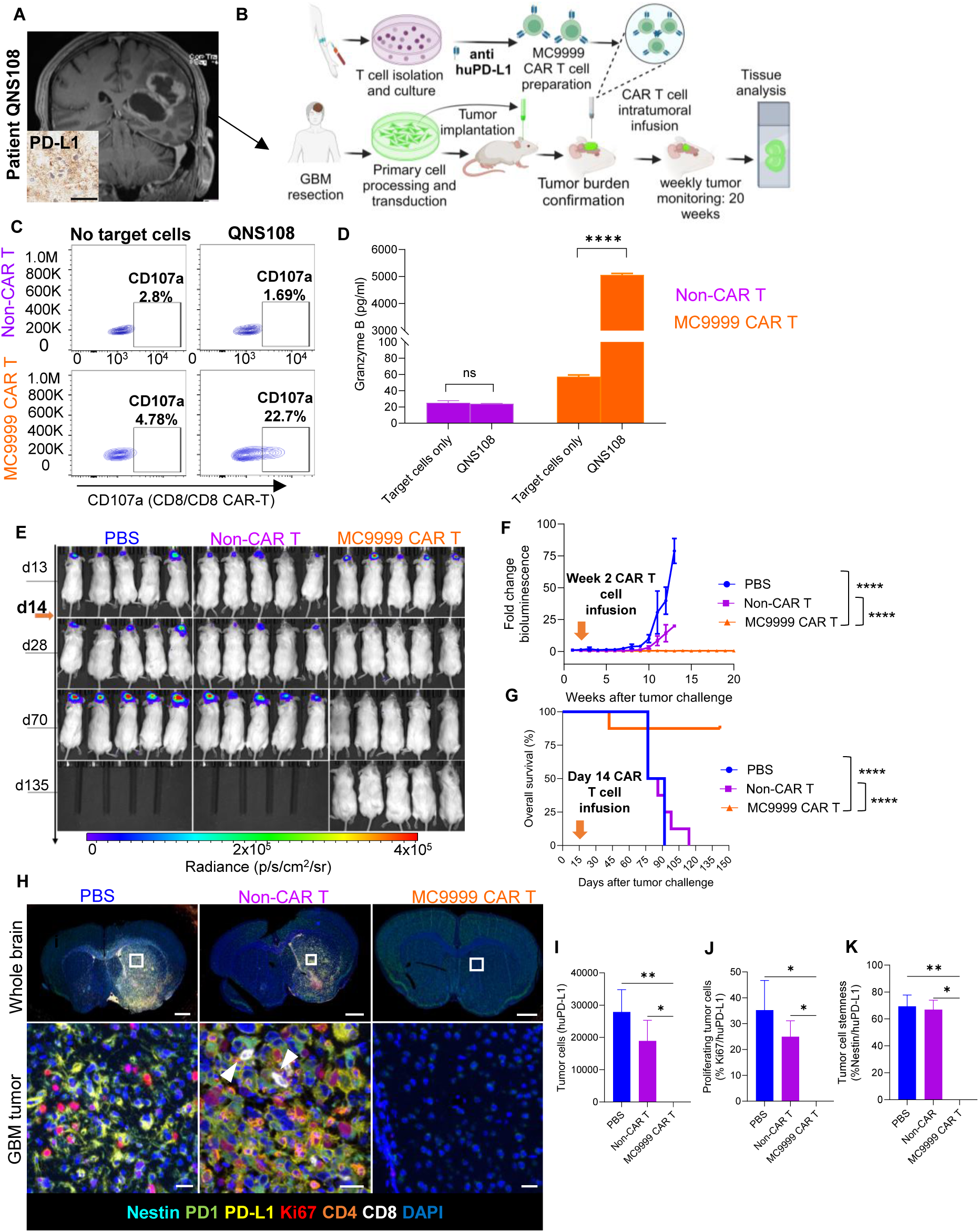
Intracranial delivery of MC9999 CAR T cells exhibits high efficiency in GBM patient derived tumor models in vitro and in vivo. **A.** T1WI Magnetic Resonance Image (MRI) of patient QNS108 showing the GBM and PD-L1 expression in tumor tissue. **B.** A schematic representation of the experimental design shows our use of healthy donor T cells for CAR T production and patient derived GBM (QNS108) engineered to develop an orthotopic xenograft model for intracranial therapeutic treatment. **C**. MC9999 CAR T cells, but not Non-CAR T cells, co-cultured QNS108 with show CD107a membrane expression. This activation was further validated by **D.** Granzyme B release (n=3; P<0.0001). **E.** MC9999 CAR T cells efficiently eradicated patient derived tumor xenografts (QNS108) in vivo. Intracranial CAR T cell infusion was performed on day 14 (orange arrow) after tumor burden confirmation. **F.** Tumors were significantly reduced in the CAR T group compared to Non-CAR T and PBS controls (n=8 per group; P<0.0001) using bioluminescence fold change. **G.** Survival was also significantly increased in the CAR T treated group (n=8 per group; P<0.0001). **H.** Analysis of the residual tumor cell population (N=5 per group) showed PD-L1, Nestin and Ki67 expression in tumor cells in the control groups but not in CAR T treated animals. Quantification shows significant increase in **I.** tumor cells (PD-L1) (Kruskall-Wallis P<0.01 for PBS vs MC9999 CAR T; P<0.05 for Non-CAR T vs MC9999 CAR T), **J.** proliferation (Ki67) (Kruskall-Wallis P<0.05 for PBS vs MC9999 CAR T; P<0.05 for Non-CAR T vs MC9999 CAR T), and **K.** stemness (Nestin) (Kruskall-Wallis P<0.01 for PBS vs MC9999 CAR T; P <0.05 for Non-CAR T vs MC9999 CAR T)in control groups compared to MC9999 CAR T group. Scale bars: 1 mm, magnification 50 µm.

To assess MC9999 CAR T cells activation by QNS108 GBM cells we performed a CD107a degranulation was measured and showed a 14-fold difference (22.7% CD107a-positive cells in CAR T group compared to1.69% in Non-CAR T group, N=3) **(Figure 2C)**. This activation was confirmed by a significant increase in granzyme B release from MC9999 CAR T cells upon incubation with QNS108 GBM cells (N=3, P<0.0001) **(Figure 2D)**.

Following this *in vitro* validation, we tested the anti-tumor efficacy *in vivo* using an orthotopic GBM xenograft model with patient-derived QNS108-PD-L1-GFP-Luc cells implanted in NSG mice brains. Fourteen days post-tumor implantation and after confirmation of tumor burden, the animals were treated with a single intracranial injection of MC9999 CAR T cells, non-CAR T cells from the same donor (allogeneic control), or PBS as a vehicle control (N=8 per group). Remarkably, after a single CAR T infusion tumors were completely eradicated within two weeks, with no recurrence detected in the 135-day observation period. In contrast, tumor size continued to grow significantly in both control groups (Non-CAR T and PBS) (N=8 per group, P<0.0001) until reaching the humane endpoint **(Figure 2E, F)**. significantly prolonged survival was observed in the CAR T-treated group compared to the control groups (N=8 per group, P<0.0001) **(Figure 2G).**

Mice brains were harvested at experimental endpoint, we assessed antigen persistence by measuring human PD-L1 expression in all groups. Tumor cells in the MC9999 CAR T-treated group were significantly reduced compared to both control groups (N=5 per group, P<0.01 for CAR T vs. PBS, P <0.05 for CAR T vs. non-CAR T) **(Figure 2H, I; Figure S2)**. Additionally, the proliferation (**Figure 2J**, N=5 per group, P <0.05 for CAR T vs. PBS, P <0.05 for CAR T vs. non-CAR T) and stemness (**Figure 2K**, N=5 per group, P <0.01 for CAR T vs. PBS, P <0.05 for CAR T vs. non-CAR T) of tumor cells were significantly reduced in the brains of mice that received MC9999 CAR T-treatment. These findings emphasize the potential of MC9999 CAR T cells as a promising therapeutic option for patient-derived GBM models.

### scRNA-seq reveals that MC9999 CD4+ and CD8+ CAR T cells elicit an IFN mediated apoptosis in patient-derived GBM cells in vivo

To gain insights into the molecular pathways driving the robust response of MC9999 CAR T cells against GBM, we performed scRNA-seq on an orthotopic xenograft model using LN229 PD-L1-OE cells. After a single intratumoral infusion of MC9999 CAR T cells or non-CAR T cells (control) (N=4 mice per group), we sacrificed animals 24 hours post-infusion, to capture both tumor and T cells, and conducted scRNA-seq **(Figure 3A)**.

**Figure 3.**
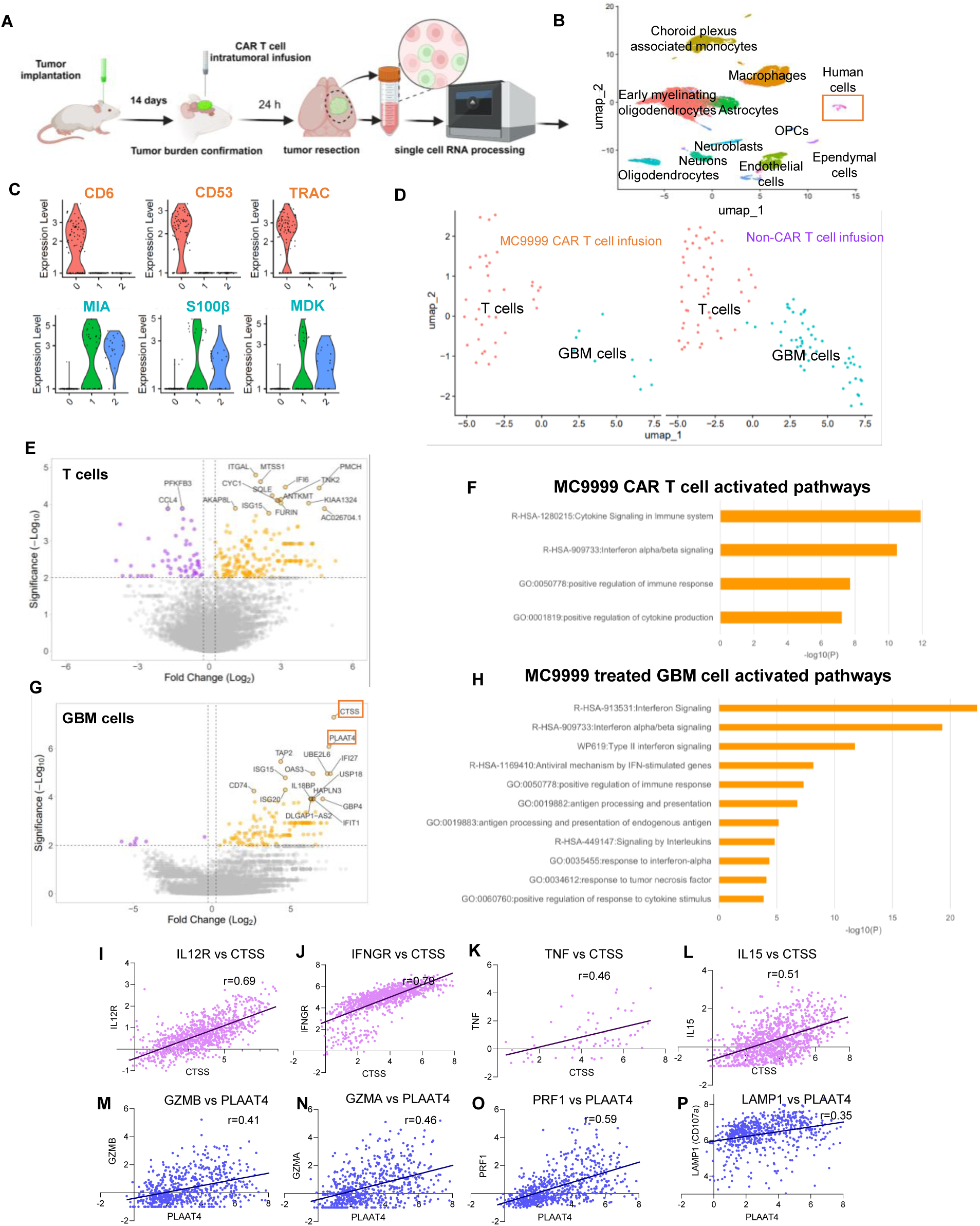
Single-cell RNA sequencing (scRNA-seq) reveals that MC 9999 CAR T treatment activates interferon-related pathways in GBM models *in vivo*. **A.** Schematic representation of the experimental approach depicting CAR T cell intratumoral infusion (N= 4 per group) after tumor burden confirmation (14 days). Animals were sacrificed 24 hours after treatment and tissue was harvested for scRNA-seq. **B.** Analysis revealed twelve distinct clusters including eleven mice brain populations and one human cluster comprised of T cells and GBM cells (orange square). **C.** Human cell cluster was reclassified into two subclusters: immune cell markers (CD6, CD53 and TRAC) shown expression in cluster 0, whereas GBM markers (MIA, S100B and MDK) were expressed in clusters 1 and 2 **D.** Re-clustering showed less tumor cells in MC9999 CAR T treated animals compared to Non-CAR T control. **E.** Differentially Expressed Gene (DEG) analysis identified upregulated genes (orange) and downregulated genes(purple) in CAR T vs. Non-CAR T cells. **F.** Pathway analysis comparing CAR T to Non-CAR T cells displayed activation of cytokine and interferon mediated cytotoxicity in T cells. **G.** CTSS and PLAAT4 (highlighted in orange rectangles) were observed to be the most upregulated genes in GBM cells. **H.** GBM cells activated pathways in response to interferon signaling and antigen presentation. **I-L**. Validation of the most upregulated genes(CTSS and PLAAT4) in GBM in response to CAR T treatment, using CGGA database, showed a significant correlation with genes related to CD4+ Th1 including **I.** IL12R (N=1011 glioma patients; R=0.62); **J.** IFNGR (N= 1018 glioma patients; R=0.79); **K.** TNF (N= 66 glioma patients; R=0.46); **L.** IL15 (N=1010 glioma patients; R=0.51) and the top upregulated gene, CTSS. **M-P**. Similarly, CD8+ cytotoxicity-related genes including **M.** GZMB (N= 649 glioma patients; R=0.41); **N.** GZMA (N= 651 glioma patients; R=0.46); **O.** PRF1 (N= 646 glioma patients; R=0.51); **P.** LAMP1 (N=651 glioma patients; R=0.35) correlated strongly with PLAAT4, the second most upregulated gene.

Twelve distinct populations were identified: 11/12 clusters represented mouse brain cells, with one smaller cluster of human GBM and T cells **(Figure 3B)**. Using markers specific to human immune cells (*CD6, CD53, CD38*) and GBM cells (*MIA, S100B, MDK*), we separated this cluster into two subpopulations **(Figure 3C)**. at this early time point, the number of GBM cells, but not T cells, were already reduced in the MC9999 CAR T-treated group **(Figure S3A-B)**. despite low numbers of cells (10 GBM cells from the MC9999 CAR T group, 52 GBM cells from the non-CAR T group, 34 MC9999 CAR T cells, and 49 non-CAR T cells **(Figure 3D)**.), differential gene expression analysis revealed numerous genes upregulated and downregulated in MC9999 CAR T cells compared to non-CAR T cells (189 upregulated DEGs and 56 downregulated DEGs) **(Figure 3E)**. Pathway analysis showed that the upregulated genes were predominantly associated with cytokine and interferon α/β signaling pathways **(Figure 3F)**. CD4+ T cells appeared as the primary activators of these pathways **(Figure S4E-G)**). Downregulated pathways (those upregulated in non-CAR T cells or downregulated in CAR T cells) in CD4+ T cells were related to migration and anaerobic metabolism, processes typically associated with T cell exhaustion ^<u>41–45</u>^ **(Figure S3F).** There were far fewer changes in CD8+ T cells which reflected activation of pathways related to interleukin signaling and transmembrane transport, suggesting roles in degranulation and cytotoxic activity **(Figure S3D,G)**. Interestingly, in GBM cells treated with MC9999 CAR T cells, DEG analysis revealed activation of interferon and TNF signaling pathways, particularly type II interferon (α/β from CD4+ T cells and γ from CD8+ T cells) and antigen processing **(Figure 3G, H)**. These results suggest the GBM cells initially responded to T cells by activating genes involved in immune signaling, which likely led to their later demise.

### Single cell-RNA seq data validation in datasets from glioma patients

To validate these findings in human glioma, we utilized data from the CGGA database to correlate the top two upregulated genes in GBM following MC9999 CAR T treatment with genes exclusively expressed by CD4+ Th1 and CD8+ T cells, both of which are known for their tumor-killing capabilities. First, we correlated the expression of *CTSS* (Cathepsin S) in GBM cells with the expression of active CD4+ Th1 markers, including *IL12R* (R=0.69, **Figure 3I**), IFNGR (R=0.79, **Figure 3J**), *TNF* (R=0.46, **Figure 3K**), and *IL15* (R=0.51, **Figure 3L**). Next, we correlated the expression of (Phospholipase A and Acyltransferase 4) *PLAAT4* with CD8+ T cell activation genes, including *GZMB* (R=0.41, **Figure 3M**), *GZMA* (R=0.46, **Figure 3N**), *PRF1* (R=0.46, **Figure 3O**), and *LAMP1* (R=0.35, **Figure 3P**). We also performed additional correlations between active CD4+ Th1 genes and *PLAAT4* **(Figure S4A-D),** showing strong correlation, while CD8+ activation genes did not correlate with *CTSS* **(Figure S4E-H).** These correlations suggest a similar mechanism of response in human glioma cells when in contact with active T cells.

Importantly, both *CTSS* and *PLAAT4* demonstrated higher expression in HGG compared to lower-grade gliomas **(Figure S4I, J),** further underscoring their relevance in the GBM response to MC9999 CAR T cell therapy. These results highlight the activation of immune-related pathways in GBM cells under attack by MC9999 CAR T cells, which is critical for their eradication.

### GBM patient derived Anti-PD-L1 (MC9999 CAR-T) cells exhibit robust cytotoxicity when incubated with their autologous tumor cells

Given that local and systemic T cells are typically dysfunctional in glioma^<u>46–48</u>^, and t that CAR T cell therapies for intracranial tumors require autologous T cells, we assessed the cytotoxic efficacy of MC9999 CAR T cells generated from GBM patient-derived T cells against their autologous BTICs. For this, we isolated both BTICs and T cells from two GBM patients (QNS985 and QNS986) (**Figure 4A, Supplementary table 2**). CAR T cells and Non-CAR T cells were manufactured from their peripheral blood, with both products passing quality control analysis **(Supplementary table 1)**. Following the same protocol, autologous BTICs were isolated as target cells for the function assessment of patient-derived MC9999 CAR T cells. IHC and flow cytometry staining confirmed positive PD-L1 expression in these BTICs **(Figure 4A and Figure S1B).**

**Figure 4.**
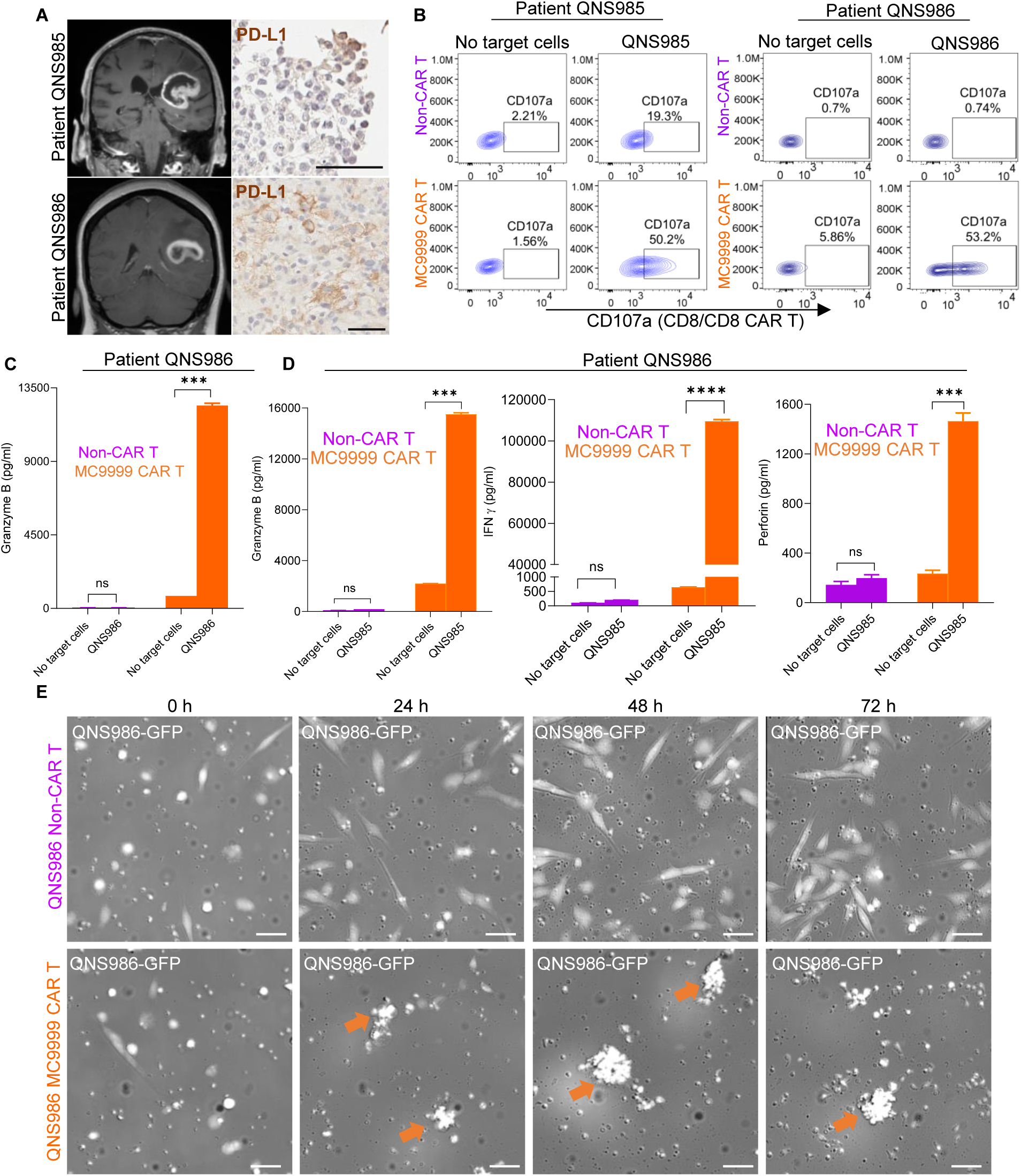
Patient derived CAR T cells show robust activation in response to the presence of their autologous tumor cells. **A.** T1WI Magnetic Resonance Image (MRI) of patients QNS985 and QNS986 showing GBMs and PD-L1 expression in tumor cells. **B-F.**, Brain tumor initiating cells (BTICs) from these patients were isolated and co-cultured with their corresponding autologous MC9999 CAR Tand Non-CAR T cells (control) to assess cytotoxicity **B.** Activation of autologous CAR T cells by autologous CAR T cells by autologous BTICs was observed, as indicated by increased CD107a expression in the CAR T group in both patients. **C.** Granzyme B release was significantly elevated in the CAR T group for both patients (N=3 per patient; P<0.001). **D.** IFNγ secretion and **E** Perforin release from patient QNS985 showed statistically significant increases in the CAR-T group (N=3; P<0.0001 and P<0.001, respectively). **F.** Representative images of CAR T cells co-cultured with autologous QNS986 tumor cells over time. Apoptotic tumor cells (orange arrows) became visibly apparent 24 hours after co-culture in the CAR T group, but not in the non-CAR T control group. Scale bars: 100 µm.

The degranulation assay showed a significant increase in CD107a membrane expression on MC9999 CAR T cells in response to autologous BTICs compared to Non-CAR T cells, with a two-fold CD107 increased in QNS985 and a 70-fold increase in QNS986 **(Figure 4B)**. This activation was further validated by elevated granzyme B release from the MC9999 CAR T-treated cells in both patient samples **(Figure 4C-D)** (N=3, P<0.001). Additionally, patient QNS985 showed significant increases in IFN-γ (**Figure 4D**, P<0.0001) and perforin A (**Figure 4D**, P<0.001) in cytokine release analysis.

To further evaluate the direct cytotoxic effects of patient-derived CAR T cells on autologous BTICs, time-lapse acquisition and analysis were conducted. BTICs from patient QNS986 were co-incubated with MC9999 CAR T or non-CAR T cells for 72 hours, capturing real-time imaging of cell proliferation and death **(Figure 4E)**. Apoptotic bodies were observed in GBM cells treated with MC9999 CAR T cells as early as six hours after treatment, while tumor cells in the Non-CAR T group continued to grow without any detected apoptotic bodies (**Figure 4E, Figure S5**).

These findings demonstrate that even T cells from GBM patients, which may have impaired functionality as patients have cancer and are on high steroid dosages that are known to impair the immune system^<u>46–48</u>^, can be engineered into potent MC9999 CAR T cells. More importantly, these CAR T cells exhibit strong cytotoxic activity against autologous tumor cells, highlighting their potential for effective personalized glioma therapy.

### GBM patient derived Anti-PD-L1 (MC9999 CAR-T) cells effectively target autologous immune suppressive populations in the tumor microenvironment

After demonstrating the direct anti-tumor efficacy of MC9999 CAR-T derived from both healthy donors and GBM patients targeting GBM tumor cell lines and autologous primary tumor BTICs, further studies were conducted to evaluate their efficacy on immunosuppressive cells within the TME. Immunosuppressive macrophages cause T cell inactivation via PD-L1 in cancer. Specifically, autologous M2 phenotype macrophages derived from GBM patients QNS924, QNS1065 and QNS1070 **(Table S2)**, as well as TAMs that were isolated from the tumor tissues of patients QNS1065 and QNS1070, were utilized to investigate the efficacy of autologous patient derived MC9999 CAR T cells on immunosuppressive cells within the TME.

CAR-T products generated from GBM patients QNS924, QNS1065 and QNS1070 achieved the quality control standards **(Table S1)**. Immunophenotyping confirmed that the target macrophages were CD163 and CD209 double-positive with PD-L1 expression **(Figure S6)**. Cytotoxicicity of the MC9999 CAR T cells was demonstrated using a degranulation assay showing significant increases in CD107a expression in MC9999 CAR-T cells against M2 macrophages—seven-fold increase in in QNS924 and a six-fold increase% in QNS1065 compared to Non-CAR T groups **(Figure 5A)**. Additionally, MC9999 CAR-T cells targeting M2, and TAMs induced substantial cytokine release, including Granzyme A, Granzyme B, and IFN-γ (**Figure 5B-D**, P<0.001). These findings demonstrate that MC9999 CAR T cells derived from patients with brain cancer can effectively target not only GBM cells but also the immunosuppressive components of the TME, enhancing their therapeutic potential.

**Figure 5.**
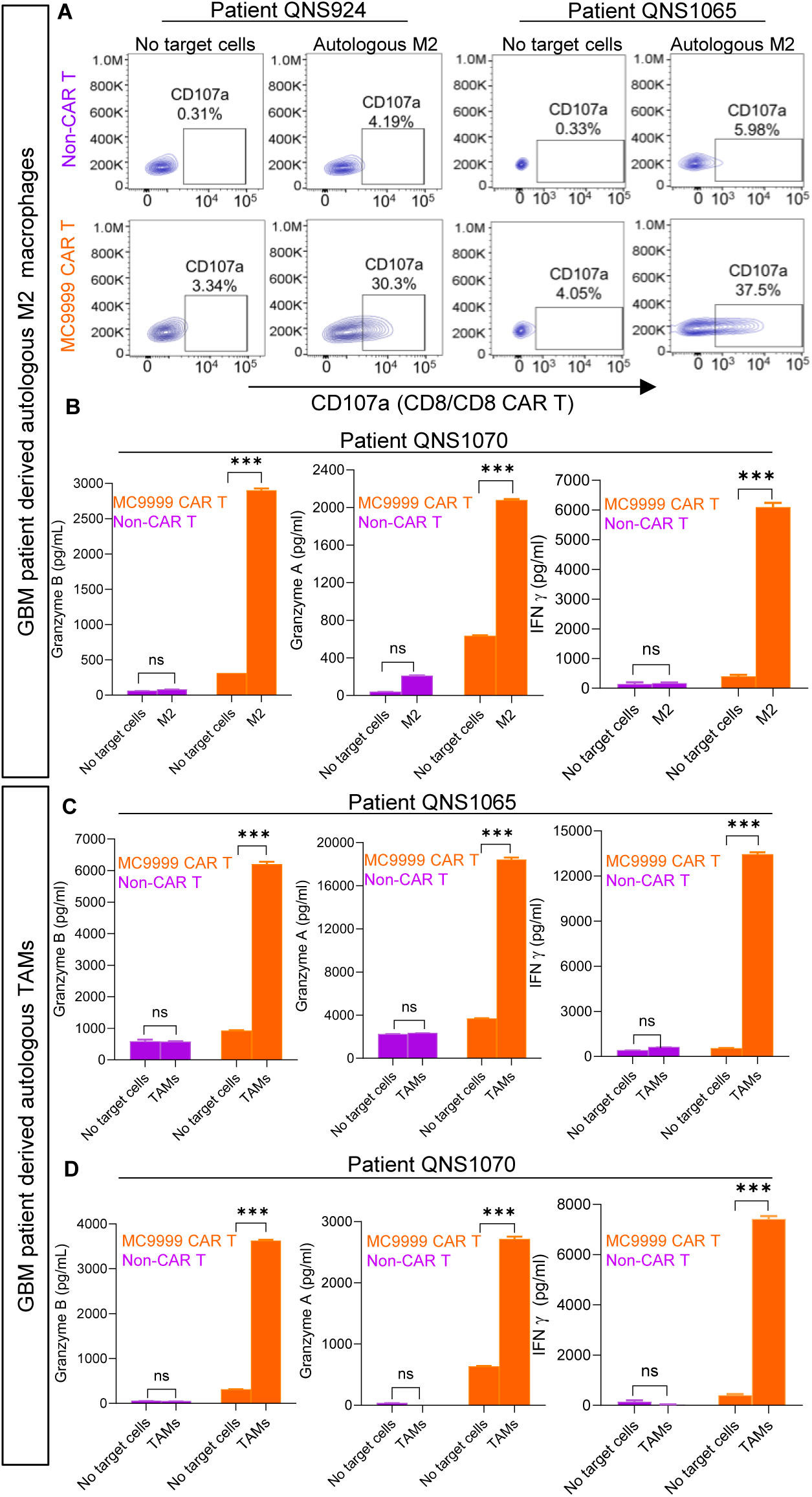
Patient derived MC9999 CAR T cells effectively target autologous M2 macrophages and TAMs. Monocyte-derived M2 macrophages from patient blood samples (N=3; patients QNS 924, QNS 1065, and QNS 1070) or TAMs isolated from tumor tissues (N=2; patients QNS 1065 and QNS 1070) were incubated with autologous MC9999 CAR T cells to assess cytotoxicity. **A.** CD107a expression increased in CD8+ CAR T cells for the two patients. **B.** Granzyme B, granzyme A, and IFNγ release were significantly elevated in CAR T group on M2 for patient QNS1070 (N=3 per experiment; P<0.001). **C.** Granzyme B, granzyme A, and IFNγ release were significantly elevated in CAR T group on autologous TAMs from patients QNS1065 and 1070 (N=3 per experiment; P<0.001)

### PD-L1 expression on high-grade glioma patient tissue and cytotoxic effects of astrocytoma patient-derived PD-L1 CAR T cells

To assess the safety of anti-PD-L1 CAR T cells for intracranial delivery in humans, we performed co-staining for neurons, glia (astrocytes and oligodendrocytes), tumor cells, blood vessels, and microglia markers with PD-L1 **(Figure 6A, Supplementary table 2)**. The results showed that PD-L1 expression was restricted to Nestin/GFAP-positive tumor cells and Iba1-positive macrophages/microglia within the tumor, with no expression detected in neurons, astrocytes, oligodendrocytes, or blood vessels. This indicates that anti-PD-L1 CAR T therapy could be safely delivered to the brain without affecting normal cell populations.

**Figure 6.**
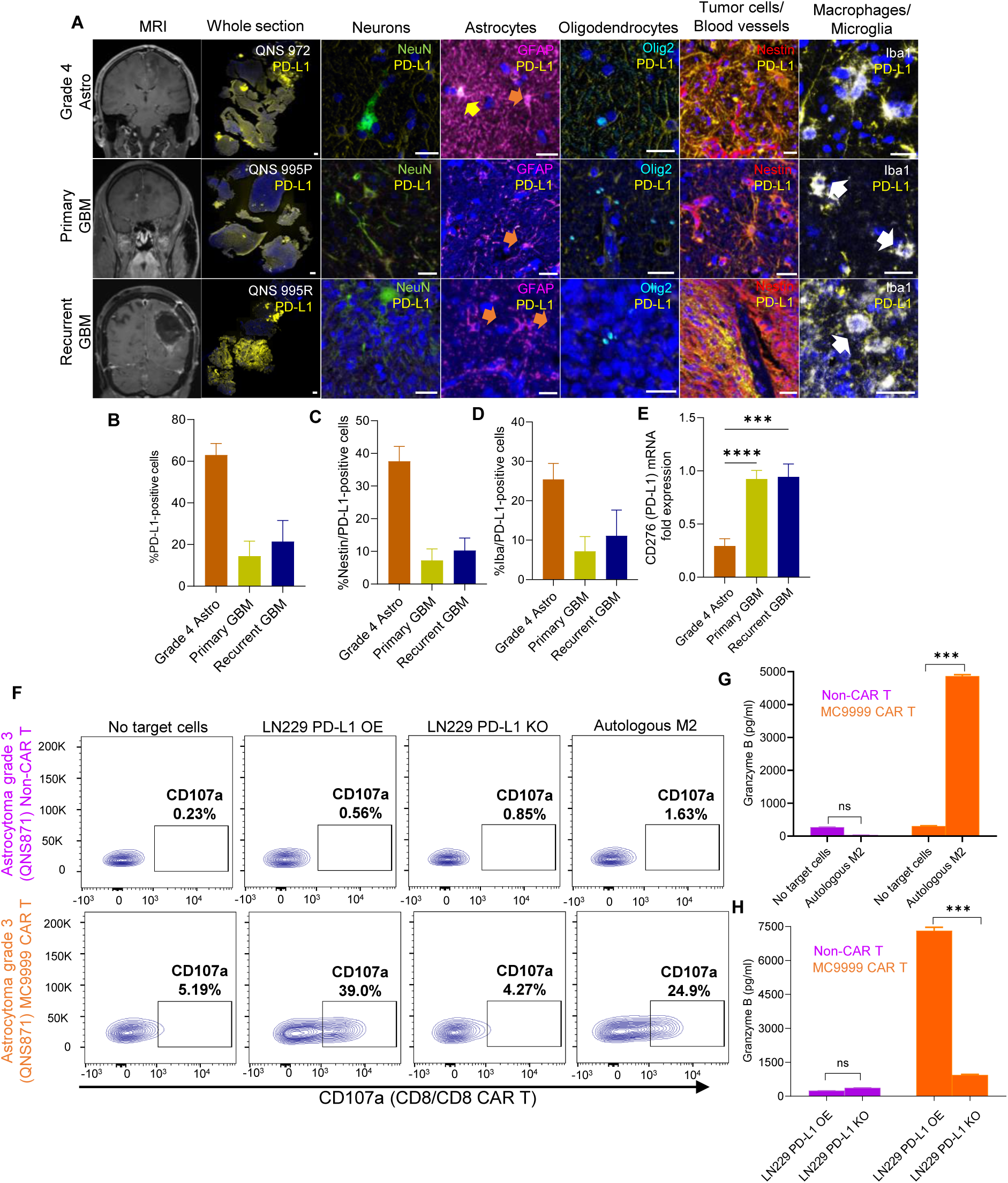
PD-L1 is a safe and effective target for anti-PD-L1 CAR T intracranial delivery and cytotoxic effects of astrocytoma patient-derived PD-L1 CAR T cells on autologous M2 macrophages. **A.** Representative mIHC staining of high-grade glioma patents including Grade 4 astrocytoma, primary and matched recurrent GBM show PD-L1 expression in tumor cells and TAMs (white arrows). Some GFAP+ cells (tumor cells) express PD-L1 (yellow arrow). Most astrocytes did not express PD-L1 (orange arrows). No Neurons (NeuN+ cells) or oligodendrocytes (Olig2+) cells co-express PD-L1. **B.** Quantification of PD-L1 in high grade glioma patients (N=4 per condition) showed all these tumors present PD-L1 in **C.** tumor cells and **D.** TAMs. **E.** Quantification and validation of PD-L1-positive cells and mRNA levels in a larger cohort (CGGA) indicate significantly higher PD-L1 mRNA in primary and recurrent glioblastomas compared to grade 4 astrocytoma (N=598; P<0.001, P<0.0001). **F-H.** Cytotoxicity of astrocytoma patient-derived PD-L1 CAR T cells on autologous M2 macrophages confirmed by degranulation and granzyme B ELISA assays. Scale bars: whole section: 1 mm, magnification: 25 µm.

Our cohort comprised 13 patient samples, including 4 matched primary and recurrent GBM cases and 5 grade 4 astrocytomas (AS). PD-L1 expression was notably high across all glioma samples **(Figure 6B)**, expressed on 14.45% in primary GBM, 21.42% in recurrent GBM, and 63.02% in grade 4 AS of the tissue analyzed. PD-L1cells co-localized with Nestin **(Figure 6C)**: 7.25% in primary GBM, 10.29% in recurrent GBM, and 37.59% in grade 4 AS, while the remainder co-expressed Iba1 **(Figure 6D)**: 7.2% in primary GBM, 11.13% in recurrent GBM, and 25.42% in grade 4 AS. Further analysis of CGGA data confirmed significantly higher PD-L1 mRNA expression was found in GBM and grade 4 AS **(Figure 6E)** (P for grade 4 AS vs. primary GBM <0.0001, P for grade 4 AS vs. recurrent GBM <0.001), supporting PD-L1 as a sufficiently abundant target in glioma patients.

To evaluate long-term safety, we analyzed mice tissue bearing QNS108 patient-derived tumors at the experimental endpoint. CD4+ T cells were observed in the meninges of CAR T-treated mice even 5 months after treatment, without any histologically detectable damage to the brain **(Figure S7A).** Staining revealed glial scar formation with abundant astrocytes and invariable number of macrophages in the treated hemisphere vs the contralateral hemisphere (Figure 7B-D), indicating effective tumor resolution with normal brain repair. Single-cell data from early stages further supported the tumor eradication **(Figure S7E)**. In MC9999 CAR T-treated animals, early myelinating oligodendrocytes and oligodendrocyte precursor cells (OPCs) were among the most abundant populations, activating pathways (i.e., regulation of synapse organization, regulation of transmembrane protein, monocarboxylic acid biosynthesis process, etc.) related to regeneration **(Figure S7E-F)**. These results demonstrate that MC9999 CAR T therapy is not only safe for intracranial use in our preclinical model but also supports normal brain healing following tumor eradication. These results demonstrate that MC9999 CAR T therapy is not only safe for intracranial use in our preclinical model but also supports normal brain healing following tumor eradication.

To further test the efficacy of our CAR T product in different glioma types, we generated MC9999 CAR T cells from a grade 3 AS patient (QNS871, **Supplementary table 2)** High PD-L1 expression of autologous M2 **(Figure S6)** served as target immunosuppressive cell models. Given the unavailability of autologous tumor cells, we used LN229 PD-L1 OE and KO cells as target tumor cells. CAR T treatment increased 70-fold CD107a expression against LN229 PD-L1 OE and 15-fold against autologous M2 macrophages **(Figure 6F)**, this finding was supported by an increase in Granzyme B release **(Figure 6G-H)**.

## DISCUSSION

Here, we present preclinical data demonstrating efficacy and preliminary safety of a high-affinity anti-PD-L1 CAR T cell product for the treatment of gliomas, with the added capability of targeting immunosuppressive macrophages within the TME via intracranial delivery. Recent clinical trials have shown that the intracranial administration of CAR T cells, including intraventricular infusions, is safe and does not result in CAR T cell leakage outside the brain compartment^<u>29</u>^. However, despite these advances, clinical trial results thus far^<u>29,30,49,50</u>^ have not consistently achieved objective radiographic response (ORR) according to the modified Response Assessment in Neuro-Oncology (RANO) criteria ^<u>51</u>^, nor have they led to durable remissions.

One possible explanation for the partial responses observed is the immunosuppressive nature of the TME, which undermines the activity of endogenous T cells recruited during the immune response (“cold”-TME). Endogenous T cells are crucial because CAR T cells can only recognize and target cells expressing the specific antigen they are engineered for, leaving gaps in the immune response when antigen loss mechanisms occur within the tumor^<u>52,53</u>^.

MC9999 CAR T construct demonstrated high specificity and potent anti-tumor effects in both *in vitro* and *in vivo* GBM patient-derived cell line models. Importantly, this activity extends beyond GBM tumor cells, also effectively targeting autologous models of PD-L1-expressing immunosuppressive macrophages, including TAMs and M2-polarized macrophages^<u>38,54,55</u>^. The CAR T cells triggered a robust immune response, driving GBM cell apoptosis through the activation of interferon-mediated apoptotic pathways. Analysis of CD4+ and CD8+ T cells from the MC9999 CAR T treatment group revealed distinct upregulation of cytokine and interferon signaling pathways, while the GBM cells responded by activating genes involved in antigen processing and immune response. This suggests that the interaction between CAR T cells and GBM cells elicits a potent immune cascade, promoting tumor cell death. These pathways are consistent with previous studies of mesenchymal-type glioma stem cells (BTICs), which often exhibit enhanced resistance to therapy^<u>16,56</u>^.

Interestingly, the pathway analysis identified distinct roles for CD4+ and CD8+ T cell subpopulations in this immune response. CD4+ T cells were found to be primarily responsible for activating interferon signaling pathways, whereas CD8+ T cells mediated cytotoxicity through degranulation. This dual mechanism of action demonstrates the versatility of MC9999 CAR T cells in orchestrating both direct cytotoxic effects and broader immune activation, further reinforcing their potential for glioma therapy. These pathways appear to be activated in the same manner in human GBM, as shown by the correlations of the top upregulated genes and Th1 CD4+ and CD8 cells.

In a clinical context, especially for intracranial delivery, it is essential to generate CAR T cells from the patient’s own immune system to mitigate risks such as graft-versus-host disease and immune rejection, particularly within the immune-privileged environment of the brain. To assess the feasibility of this approach, we engineered autologous T cells from GBM patients to express the MC9999 CAR. These patient-derived CAR T cells exhibited strong cytotoxicity against their respective BTICs, echoing the potency seen with healthy donor-derived CAR T cells. CD107a expression and granzyme B release were significantly elevated in the MC9999 CAR T cells, confirming their capacity for tumor cell killing. Moreover, these CAR T cells were able to target not only GBM tumor cells but also the immunosuppressive cells within the TME, specifically TAMs and M2 macrophages, which are known to inhibit anti-tumor immune responses. By targeting these immunosuppressive cells, MC9999 CAR T cells may overcome key barriers to therapeutic efficacy, enhance the overall anti-tumor activity within the GBM microenvironment.

Patient-derived models have provided compelling evidence that MC9999 CAR T cells hold considerable therapeutic potential against GBM. By specifically targeting PD-L1, these CAR T cells effectively kill tumor cells, eradicate tumors *in vivo*, and surmount the immunosuppressive hurdles posed by the TME. As we move toward clinical translation, further work is needed to optimize delivery methods and refine strategies to maintain CAR T cell persistence and function. Nonetheless, these findings establish a strong foundation for the continued development of MC9999 CAR T cells as a promising treatment modality for GBM, offering new hope to patients.

## METHODS

### T cell isolation

For healthy volunteer donors’ peripheral blood mononuclear cells (PBMCs) were collected via leukapheresis using leukocyte reduction system (LRS) cones, by the Division of Transfusion Medicine, Mayo Clinic, Rochester, Minnesota, following current regulatory requirements^<u>57</u>^. To isolate naïve and memory T cell (Tn/mem) from PBMCs we used a three-step procedure, involving negative selection of both CD14 and CD25, followed by positive selection of CD62L, using CD14, CD25, and CD62L microbeads (Miltenyi Biotech)^<u>53</u>^. GBM patient blood was collected with informed consent and approved by the Mayo Clinic, Florida Institutional Review Board (IRB# 17-003013-32). Briefly, we first isolated PBMCs, then used Pan T cell isolation kit (Miltenyi Biotec) to isolate T cells, as previously described^<u>31</u>^.

### CAR T cell generation

To engineer MC9999 CAR T cells we used a high affinity PD-L1 scFv^<u>58</u>^, a hinge region with a CD4 transmembrane domain, as well as 4-1BB and CD3ζ intracellular signaling domains, complemented with a truncated EGFR^<u>31</u>^. The CAR cDNA was integrated into pHIV.7 lentiviral vector. To ensure efficient lentivirus production, we utilized 293FT cells, followed by concentration and titer determination using Jurkat cells. Tn/mem or Pan-T cell populations, isolated from PBMCs, were divided into two aliquots for generating Non-CAR T-cells as control and another for generating CAR T-cells followed the previous protocols^<u>53</u>^.

### GBM cell isolation and engineering

Patient tumor tissue was collected with informed consent and approved by the Mayo Clinic, Florida Institutional Review Board (IRB# 16-008485)^<u>59</u>^ and Brain Tumor Initiating Cells (BTICs) from patients were generated as previously described^<u>31,60</u>^. To engineer LN229, QNS 108, and QNS 986), we transduced them with GFP luciferase (GFP-luc) and PD-L1 lentivirus, respectively. We engineered LN229 cells with PD-L1 overexpression (OE) and knockout (KO) variants, following procedures detailed in^<u>31</u>^.

### Degranulation assay

Following the procedure in^<u>61</u>^, CAR T cells were incubated with target cells at a 2:1 E:T ratio in RPMI 1640 with GolgiStop™ and CD107a APC antibody for 6 hours. Cells were then stained with CD3, CD4, CD8, and EGFR antibodies and analyzed using an Attune or Fortessa flow cytometer, with non-CAR T cells as negative controls.

### Granule release ELISA assays

CAR T-cells and target cells were co-incubated for 72 hours at an E:T ratio of 4:1, the supernatant was collected and evaluated for granule protein release as described in^<u>61</u>^.

### Impedance-based tumor cell killing assay (xCELLigence)

To investigate the direct killing of CAR-T cells on tumor cells, impedance-based tumor cell killing assay were performed as previously described in^<u>31</u>^.

### Orthotopic xenograft model, intracranial CAR T infusion, and analysis

NOD SCID gamma (NSG) mice (8-12 weeks old-female for LN229 and male for QNS108 matching the sex of the patient from which we derived the cell line) received 3×10^5^ cells (LN229-PDL1-OE-GFP-luc)/5×10^5^ cells (QNS 108-PD-L1-OE-GFP-luc) intracranially in the striatum. Mice were randomized into three test groups (MC9999 CAR T, Non-CAR T, and PBS) test groups (16 mice per group in LN229, 8 mice per group in QNS 108). Two weeks after tumor implantation and after tumor burden confirmation using IVIS® bioluminescence imaging system, mice received 2×10^6^ T cells (MC9999 CAR T or non-CAR T) or PBS intratumorally. Tumor burden was quantified weekly by bioluminescent signal intensity. Animals were sacrificed by Ketamine-Xylazine overdose, followed by intracardiac infusion of 0.9% NaCl and 4% Paraformaldehyde. Brains were harvested and kept in the same fixative solution overnight, then they were rinsed with PBS and embedded in paraffin following standard histology procedures. Tumor growth was calculated by fold change in bioluminescence intensity. Survival data were presented and reported in Kaplan–Meier plots. Statistical analysis for survival was performed using Log-rank analysis (Mantel-Cox), for changes in tumor growth (bioluminescence-fold change) we used Kruskall-Wallis. All analyses were performed on PRISM-GraphPad 10). All animal procedures were approved by Mayo Clinic IACUC (IACUC# A00006674).

### Single cell RNA-sequencing in vivo experimental procedure

Briefly, we performed the same orthotopic xenograft model using LN229PD-L1-OE-GFP-luc (N=8 mice). Two weeks after tumor implantation, we confirmed tumor burden and intratumorally injected 2×10^6^ T cells (MC9999 CAR T (N=4) or Non-CAR T (N=4)) in 5 μl of PBS. 24 hours after T cell infusion, we sacrificed the animals by Ketamine-Xylazine overdose, intracardiac perfused them with 0.9% NaCl for 5 minutes, harvested brain, isolated tumor area and dissociated the tissue into single cell suspensions using Tumor Dissociation Kit (Miltenyi) and gentleMACS Octo Dissociator (Milteny Biotech) following manufacturer’s instructions. Then, we quantified the number of cells and performed 10X Genomics Chromium Next GEM Single Cell 3’ Kit v3.1 according to manufacturer’s instructions targeting a recovery of 1×10^4^ cells at a concentration of 1.2×10^3^ cells/uL).

### Single cell RNA-sequencing processing and data analysis

The single cell sequence preprocessing was performed using the standard 10X Genomics Cell Ranger Single Cell Software Suite (v.8.0.0). Briefly, raw sequencing data were demultiplexed and aligned to both the mouse transcriptome mm10 and human transcriptome GRCh38 to capture both mouse and human genes. Ambient RNA was removed in each sample using the DecontX algorithm^<u>62</u>^. For quality assurance, cells with less than 200 or more than 5000 unique genes were removed from downstream anlaysis. Cell filtration, QC and normalization were performed using standard Seurat package procedures (v.5.1.0). Cell clustering was performed using the Harmony algorithm^<u>63</u>^ and visualized using Uniform Manifold Approximation and Projection (UMAP). Cell clusters were annotated into unique cell types using classical cell type marker genes. Genes in each cell type were tested for differential expression between the CART and non-CART groups using the MAST model^<u>64</u>^. Pathway analysis of differentially expressed genes from each cell type was performed using Metascape (v3.5) ^<u>65</u>^.

### Human data validation for single cell RNA sequencing

To validate the top two upregulated genes found in the GBM cells of MC9999 CAR T treated animals (CTSS and PLAAT4) we used CGGA mRNA expression data (http://www.cgga.org.cn/) for genes related to T CD4+-Th1 cells (IL12R, IFNGR, TNF, and IL15) and genes related to T CD8+ cells (GZMB. GZMA, PRF1, LAMP1) for all types of diffuse gliomas according to 2021 WHO classification (Oligodendroglioma, Anaplastic Oligodendroglioma, Astrocytoma, Anaplastic Astrocytoma, and Glioblastoma) and correlated their expression to CTSS and PLAAT4 using linear correlation and Spearman r regression in PRISM, GraphPad 10.

### Human and rodent tissue staining and quantification

For histological tissue staining we used Akoya Opal 6-plex kit. Briefly, we dissolved the paraffin in the tissue following standard histological procedures, rinsed the tissue twice in ddH2O and fixed for 20 minutes in 3% Non-Buffered Formalin. We performed microwave antigen retrieval prior to every cycle using the AR buffer (pH 6 or 9) recommended by primary antibody manufacturer. For every cycle we blocked for non-specific binding for 10 minutes in primary diluent (Akoya), incubated the primary antibody for 1 hour, rinsed with TBS-T, incubated with HRP secondary for 10 minutes, rinsed with TBS-T, and incubated with the corresponding opal for 10 mins. For Opal 780, we incubated with DIG-TSA 1:100 for 1 hour before Opal 780 incubation. We performed three different panels. (1:150 mouse anti-Nestin (Millipore-SIGMA), 1:750 rabbit anti-PD-L1 (abcam), 1:200 rabbit anti-PD-1 (abcam), 1:500 rabbit anti CD4 (abcam), 1:150 rabbit anti-Ki67 (ThermoFisher), 1:250 rabbit anti-CD8 (abcam), 1:300 rabbit-anti Olig2 (Millipore-SIGMA), 1:200 rabbit anti-NeuN (Cell Signaling Technology), 1:500 rabbit anti-GFAP (Dako), 1:4000 rabbit anti-Iba1 (abcam), and 1:250 rabbit anti-Vimentin (abcam)). For quantification we used QuPath 0.5.1 object classification plugin. Statistical analysis was performed using % of positive cells for each marker, multiple comparisons were done using Kruskall Wallis and Dunn’s multiple comparisons in GraphPad PRISM 10.

### Timelapse acquisition and analysis

For autologous patient GBM-T cell experiment we seeded 5×10^3^ QNS986-GFP-Luc. 12 hours after, we added 2.5×10^4^ T cells (MC9999 CAR T or Non-CAR T) -E:T ratio: 5:1- and performed a timelapse experiment for 72 hours in Livecyte. Images were taken every 5 minutes. We used Livecyte analysis software to obtain the videos.

### Monocyte derived M2 macrophages from patient blood and TAMs isolation from tumor tissue

We cultured Monocyte derived M2 macrophages from GBM patient blood samples and followed the procedure as we have previously published^23^. CD14-positive monocytes were isolated from PBMCs using CD14 microbeads (Miltenyi) and differentiated into macrophages with M-CSF over 7 days. M2 polarization was induced on day 7 with Interleukin 4 and M-CSF for 48-72 hours, after which cells were harvested and analyzed for phenotype and PD-L1 expression via Fortessa flow cytometry. TAMs were isolated from GBM patient tumor tissue as previous described in^27^ and their immunophenotype and PD-L1 expression were analyzed using Fortessa flow cytometry (BD).

## Supporting information

Supplementary figure 1. PD-L1 expression in GBM cell lines. A. High levels of PD-L1 expression were observed in LN229 PD-L1 overexpressing (OE) cells

Supplementary figure 2. H&E staining of orthotopic mice models at experimental endpoint. A. Representative section of mice tumors from LN229 cell inje

Supplementary figure 3. Analysis and validation of scRNA seq. A. Number of GBM cells found 24 hours after CAR T/non-CAR T cell intratumoral infusion.

Supplementary figure 3. Analysis and validation of scRNA seq. A. Number of GBM cells found 24 hours after CAR T/non-CAR T cell intratumoral infusion.

Supplementary Figure 5. Timelapse video of CAR-T cell -mediated killing of autologous GBM cells (QNS986) A. incubated with MC9999 CAR T cells; B. Incu

Supplementary Figure 6. Immunophenotypic Characterization and PD-L1 Expression in M2 Macrophages and TAMs from GBM Patients. Macrophages were identifi

Supplementary figure 7. Analysis of brain reorganization after CAR T infusion. A. Analysis of microglia and astrocytes of PBS, Non-CAR T and MC9999 CA

Supplementary table 1: QC analysis for patient derived CAR T cells

Supplementary table 2: Patient clinical and demographic description

## ACKNOWLEDGEMENTS

A.Q.-H. was supported by the Mayo Clinic Clinician Investigator award, the William J. and Charles H. Mayo Named Professorship, the Monica Flynn Jacoby Endowed Chair, and the Uihlein Neuro-Oncology Research Fund. We would like to acknowledge the funding support to H.Q., which includes the Florida Health Grant, the Mayo Clinic Florida CAR-T Program Fundand the Florida Department of Medicine Team Science Award. Mayo Clinic President’s Discovery Translation Program Award, Florida State DeSantis Funds awarded to M.K-D and H.Q. and the generosity of benefactor funds from Lynch Family. R. R.-V was supported by Fourth and Gold Inc. M.J.U-N was supported by Mayo Clinic Center for Brain Tumor Innovation and Uilhein professorship research grant. J.E.S-G was supported by Uilhein professorship research grant. We would finally like to Acknowledge Brandi Edenfield for Histology processing and all our patients for their generosity sharing their tissue for research.

## AUTHOR CONTRIBUTIONS

M.J.U-N, Y.L., J.E.S-G., Y.Q., V.K.J, M.M.B., M.A.B, T.H., C.G-P., A.D.B., R.R-V, A.N.B., -F.Q. T-N, and J.B.H. performed the experiments M.J.U-N, Y.L, and Y.R. analyzed the experiments. M.J.U-N, Y.L, S.R.I., R.D., M.A.K-D., H.D. H.Q., and A.Q.-H, and designed the experiments. M.J..U-N., Y.L., J.E.S-G, H.Q., and A.Q-H. wrote the manuscript. All authors read, edited, and approved the article.

## DECLARATION OF INTERESTS

M.K.-D: Grant/research support: Bristol Myers Squibb, Novartis and Pharmacyclics

M.K-D: Lecture/honorarium: Kite Pharma

H.Q: is a is a founder of AquaLux and a member of its scientific advisory board

A.Q-H: is a is a founder of AquaLux and a member of its scientific advisory board

A.Q-H, H.Q., R.D.,Y.L., H.D: Declare MC9999 CAR T patent **US patent filing 68,892**, Technology= 2023-199, Case No. DR23-557

