## Supplementary figure 1. PD-L1 expression in GBM cell lines. A. High levels of PD-L1 expression were observed in LN229 PD-L1 overexpressing (OE) cells for "Single CAR-Dual target: Intracranial Delivery of Anti-PD-L1 CAR T Cells Effectively Eradicates Glioma and Immunosuppressive Cells in the Tumor Microenvironment"

### Slide 1
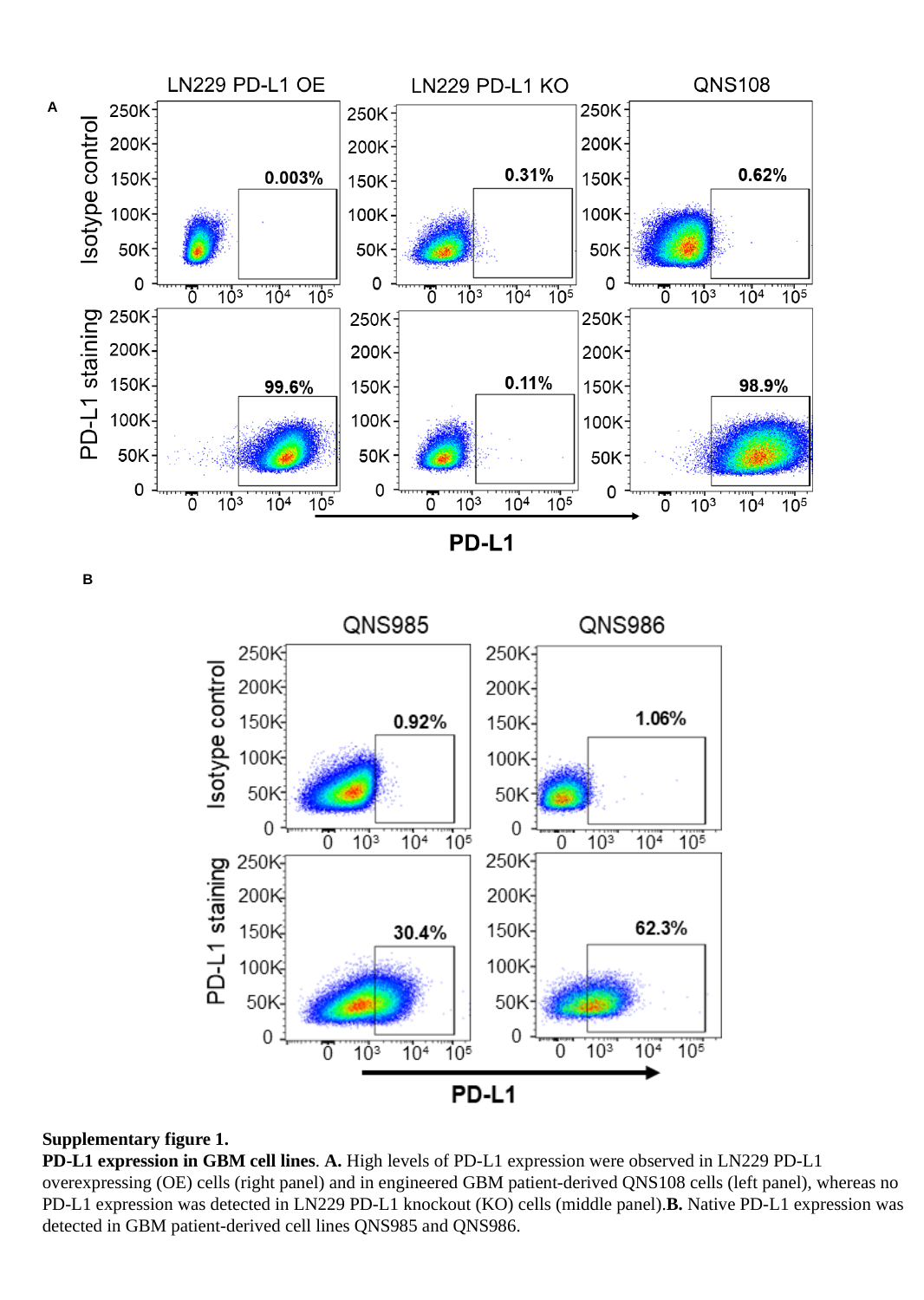

A
B
Supplementary figure 1.
PD-L1 expression in GBM cell lines. A. High levels of PD-L1 expression were observed in LN229 PD-L1 overexpressing (OE) cells (right panel) and in engineered GBM patient-derived QNS108 cells (left panel), whereas no PD-L1 expression was detected in LN229 PD-L1 knockout (KO) cells (middle panel).B. Native PD-L1 expression was detected in GBM patient-derived cell lines QNS985 and QNS986.
