## Supplementary figure 3. Analysis and validation of scRNA seq. A. Number of GBM cells found 24 hours after CAR T/non-CAR T cell intratumoral infusion. for "Single CAR-Dual target: Intracranial Delivery of Anti-PD-L1 CAR T Cells Effectively Eradicates Glioma and Immunosuppressive Cells in the Tumor Microenvironment"

### Slide 1
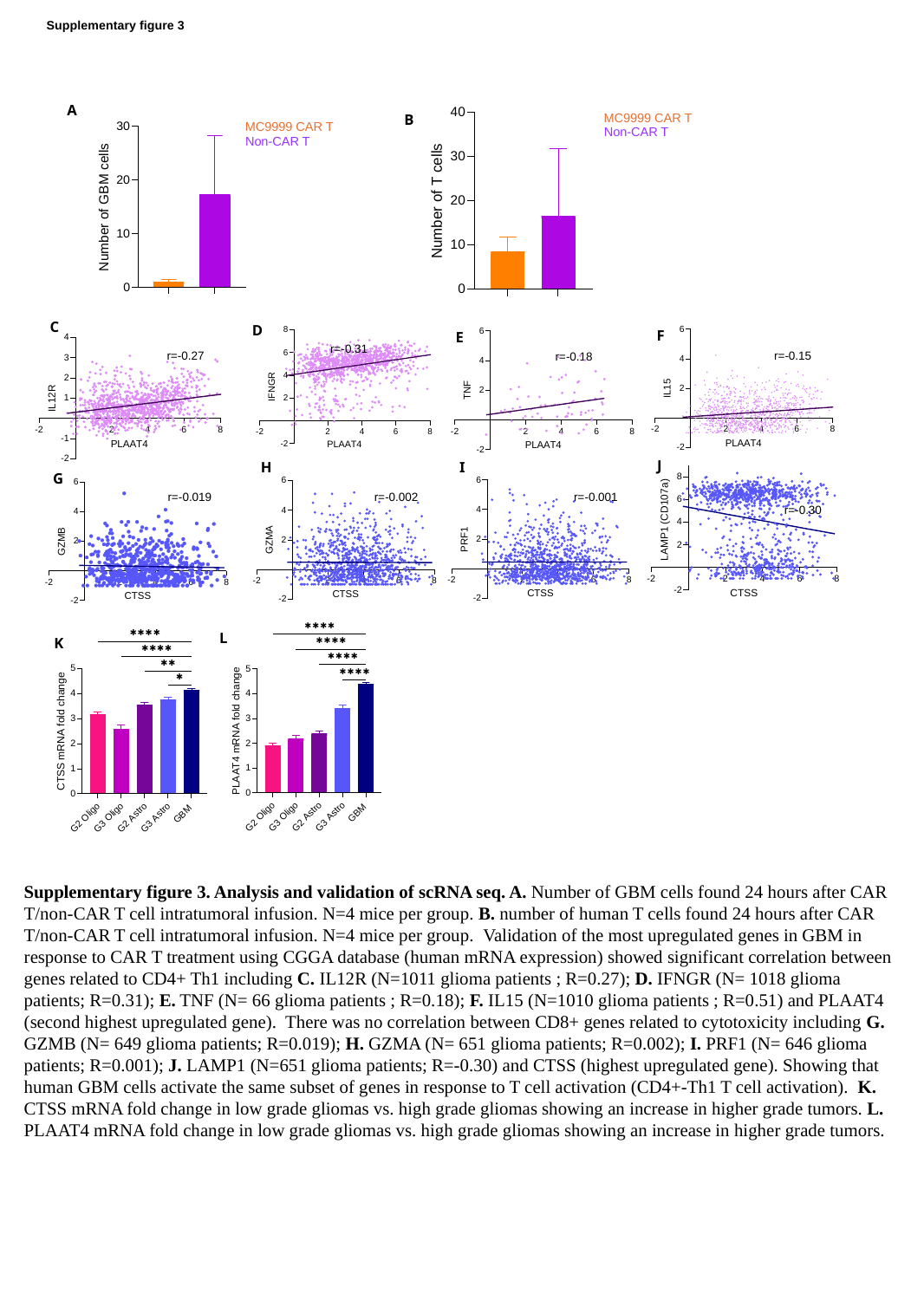

Supplementary figure 3
A
MC9999 CAR T
B
MC9999 CAR T
Non-CAR T
Non-CAR T
C
D
F
E
r=-0.31
r=-0.27
r=-0.15
r=-0.18
J
H
I
G
r=-0.002
r=-0.019
r=-0.001
r=-0.30
L
K
Supplementary figure 3. Analysis and validation of scRNA seq. A. Number of GBM cells found 24 hours after CAR T/non-CAR T cell intratumoral infusion. N=4 mice per group. B. number of human T cells found 24 hours after CAR T/non-CAR T cell intratumoral infusion. N=4 mice per group. Validation of the most upregulated genes in GBM in response to CAR T treatment using CGGA database (human mRNA expression) showed significant correlation between genes related to CD4+ Th1 including C. IL12R (N=1011 glioma patients ; R=0.27); D. IFNGR (N= 1018 glioma patients; R=0.31); E. TNF (N= 66 glioma patients ; R=0.18); F. IL15 (N=1010 glioma patients ; R=0.51) and PLAAT4 (second highest upregulated gene). There was no correlation between CD8+ genes related to cytotoxicity including G. GZMB (N= 649 glioma patients; R=0.019); H. GZMA (N= 651 glioma patients; R=0.002); I. PRF1 (N= 646 glioma patients; R=0.001); J. LAMP1 (N=651 glioma patients; R=-0.30) and CTSS (highest upregulated gene). Showing that human GBM cells activate the same subset of genes in response to T cell activation (CD4+-Th1 T cell activation). K. CTSS mRNA fold change in low grade gliomas vs. high grade gliomas showing an increase in higher grade tumors. L. PLAAT4 mRNA fold change in low grade gliomas vs. high grade gliomas showing an increase in higher grade tumors.
