## Supplementary figure 3. Analysis and validation of scRNA seq. A. Number of GBM cells found 24 hours after CAR T/non-CAR T cell intratumoral infusion. for "Single CAR-Dual target: Intracranial Delivery of Anti-PD-L1 CAR T Cells Effectively Eradicates Glioma and Immunosuppressive Cells in the Tumor Microenvironment"

### Slide 1
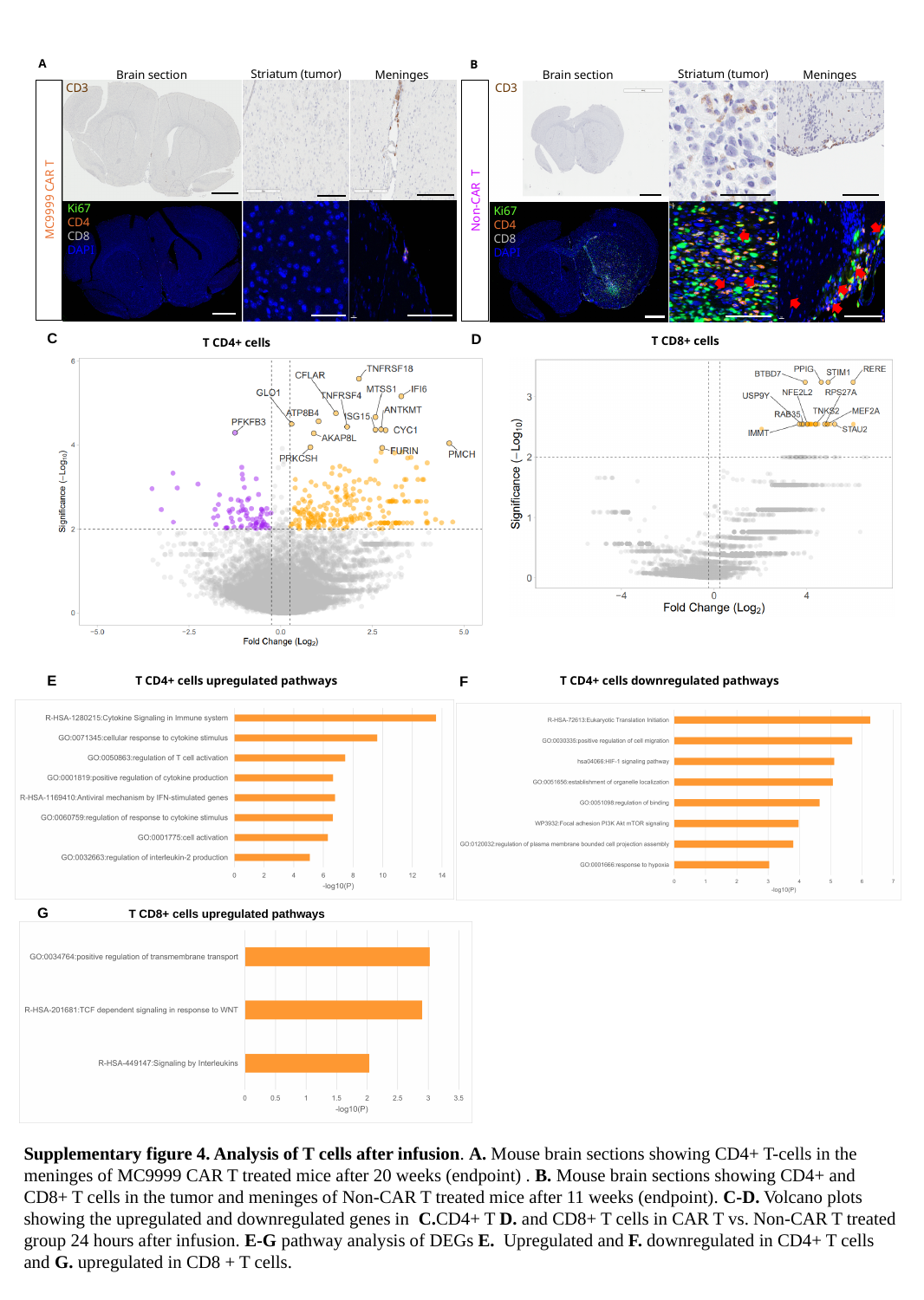

A
B
Striatum (tumor)
Striatum (tumor)
Brain section
Meninges
Brain section
Meninges
CD3
CD3
MC9999 CAR T
Non-CAR T
Ki67
CD4
CD8
DAPI
Ki67
CD4
CD8
DAPI
C
D
T CD8+ cells
T CD4+ cells
E
F
T CD4+ cells upregulated pathways
T CD4+ cells downregulated pathways
G
 T CD8+ cells upregulated pathways
Supplementary figure 4. Analysis of T cells after infusion. A. Mouse brain sections showing CD4+ T-cells in the meninges of MC9999 CAR T treated mice after 20 weeks (endpoint) . B. Mouse brain sections showing CD4+ and CD8+ T cells in the tumor and meninges of Non-CAR T treated mice after 11 weeks (endpoint). C-D. Volcano plots showing the upregulated and downregulated genes in C.CD4+ T D. and CD8+ T cells in CAR T vs. Non-CAR T treated group 24 hours after infusion. E-G pathway analysis of DEGs E. Upregulated and F. downregulated in CD4+ T cells and G. upregulated in CD8 + T cells.
