## Supplementary Figure 6. Immunophenotypic Characterization and PD-L1 Expression in M2 Macrophages and TAMs from GBM Patients. Macrophages were identifi for "Single CAR-Dual target: Intracranial Delivery of Anti-PD-L1 CAR T Cells Effectively Eradicates Glioma and Immunosuppressive Cells in the Tumor Microenvironment"

### Slide 1
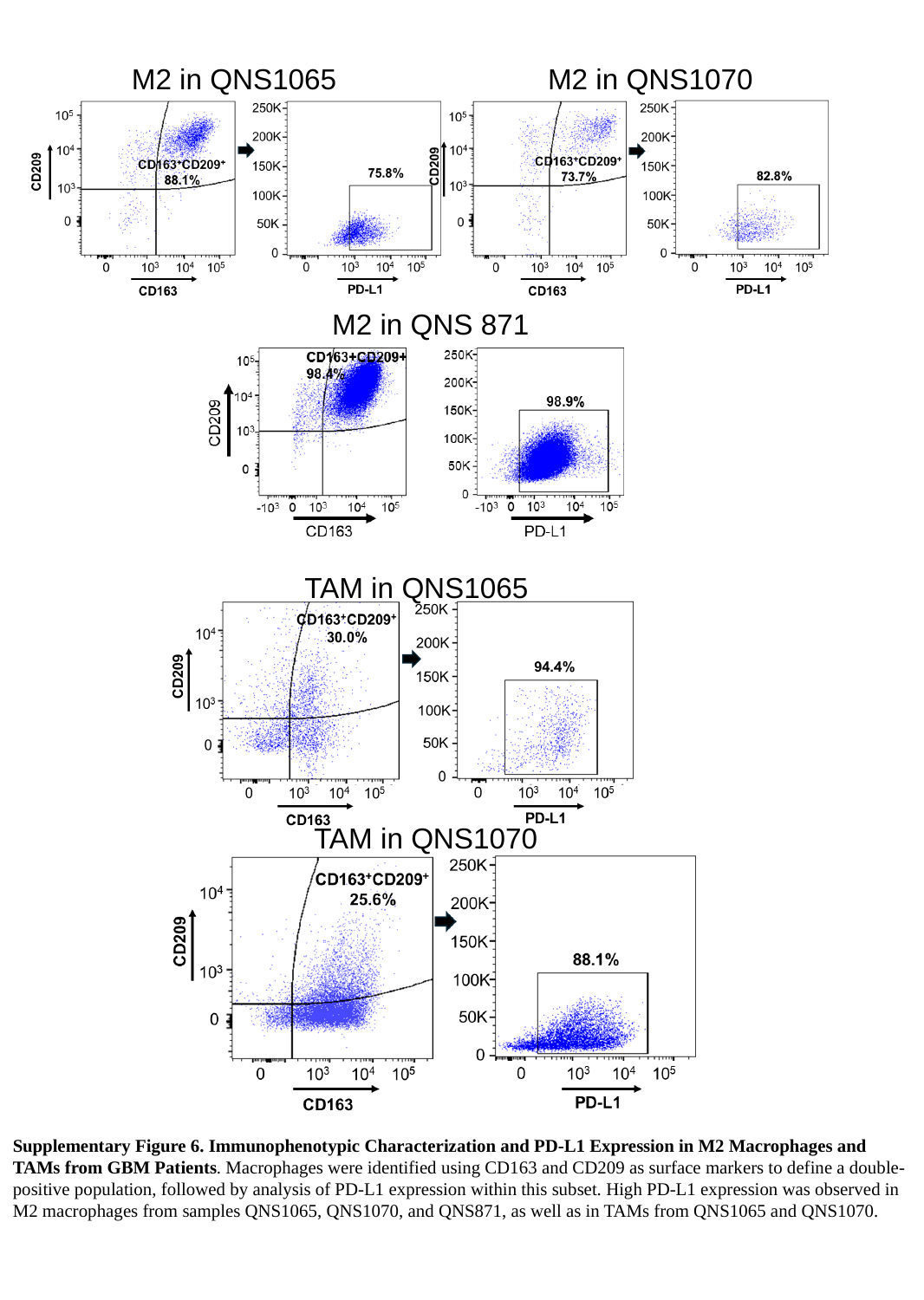

M2 in QNS1070
M2 in QNS1065
M2 in QNS 871
TAM in QNS1065
TAM in QNS1070
Supplementary Figure 6. Immunophenotypic Characterization and PD-L1 Expression in M2 Macrophages and TAMs from GBM Patients. Macrophages were identified using CD163 and CD209 as surface markers to define a double-positive population, followed by analysis of PD-L1 expression within this subset. High PD-L1 expression was observed in M2 macrophages from samples QNS1065, QNS1070, and QNS871, as well as in TAMs from QNS1065 and QNS1070.
