## Supplementary figure 7. Analysis of brain reorganization after CAR T infusion. A. Analysis of microglia and astrocytes of PBS, Non-CAR T and MC9999 CA for "Single CAR-Dual target: Intracranial Delivery of Anti-PD-L1 CAR T Cells Effectively Eradicates Glioma and Immunosuppressive Cells in the Tumor Microenvironment"

### Slide 1
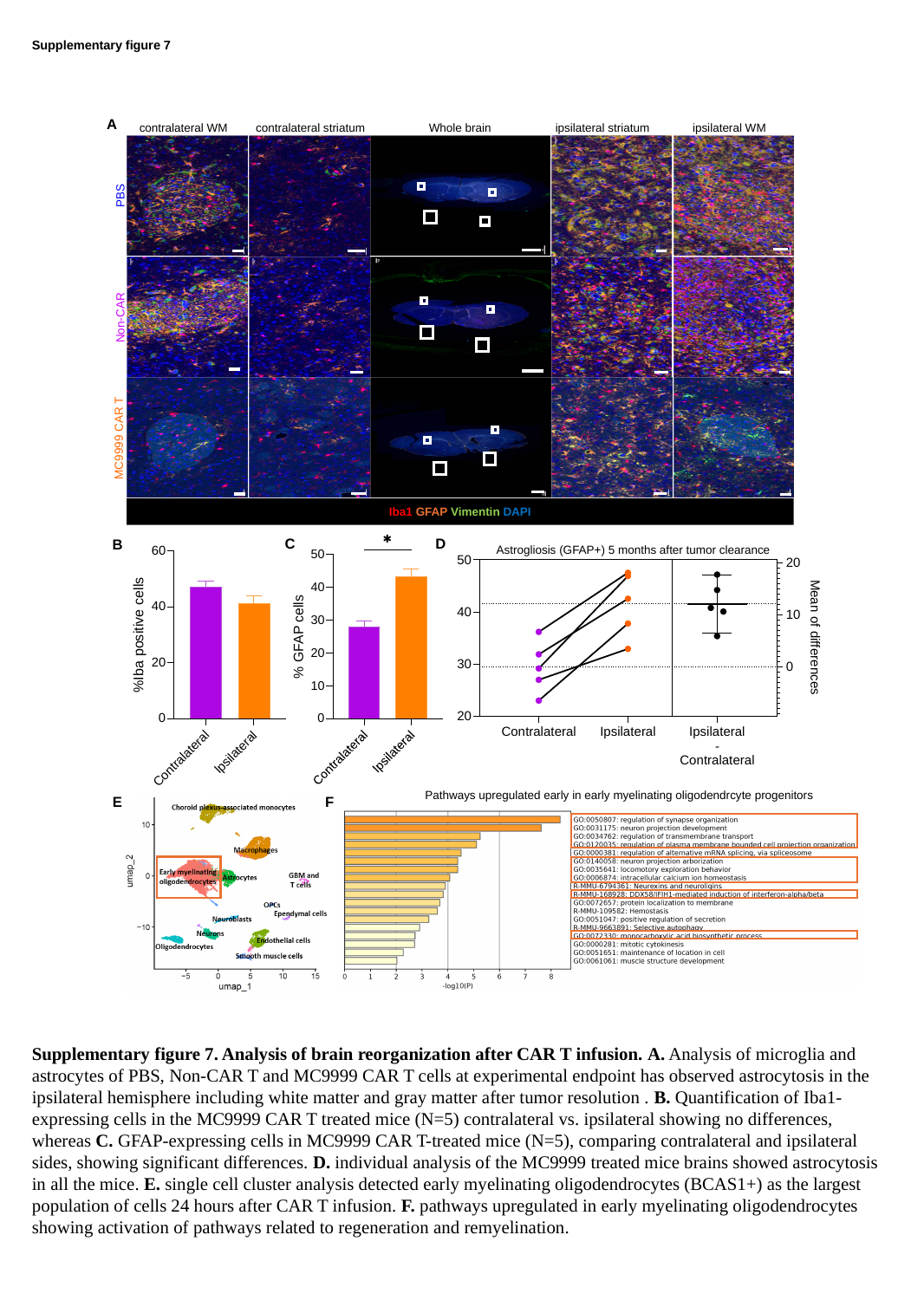

Supplementary figure 7
A
 contralateral WM
 contralateral striatum
Whole brain
 ipsilateral striatum
 ipsilateral WM
PBS
Non-CAR
MC9999 CAR T
Iba1 GFAP Vimentin DAPI
C
D
B
Astrogliosis (GFAP+) 5 months after tumor clearance
Pathways upregulated early in early myelinating oligodendrcyte progenitors
F
E
Supplementary figure 7. Analysis of brain reorganization after CAR T infusion. A. Analysis of microglia and astrocytes of PBS, Non-CAR T and MC9999 CAR T cells at experimental endpoint has observed astrocytosis in the ipsilateral hemisphere including white matter and gray matter after tumor resolution . B. Quantification of Iba1-expressing cells in the MC9999 CAR T treated mice (N=5) contralateral vs. ipsilateral showing no differences, whereas C. GFAP-expressing cells in MC9999 CAR T-treated mice (N=5), comparing contralateral and ipsilateral sides, showing significant differences. D. individual analysis of the MC9999 treated mice brains showed astrocytosis in all the mice. E. single cell cluster analysis detected early myelinating oligodendrocytes (BCAS1+) as the largest population of cells 24 hours after CAR T infusion. F. pathways upregulated in early myelinating oligodendrocytes showing activation of pathways related to regeneration and remyelination.
