## Supplementary table 1: QC analysis for patient derived CAR T cells for "Single CAR-Dual target: Intracranial Delivery of Anti-PD-L1 CAR T Cells Effectively Eradicates Glioma and Immunosuppressive Cells in the Tumor Microenvironment"

### Slide 1
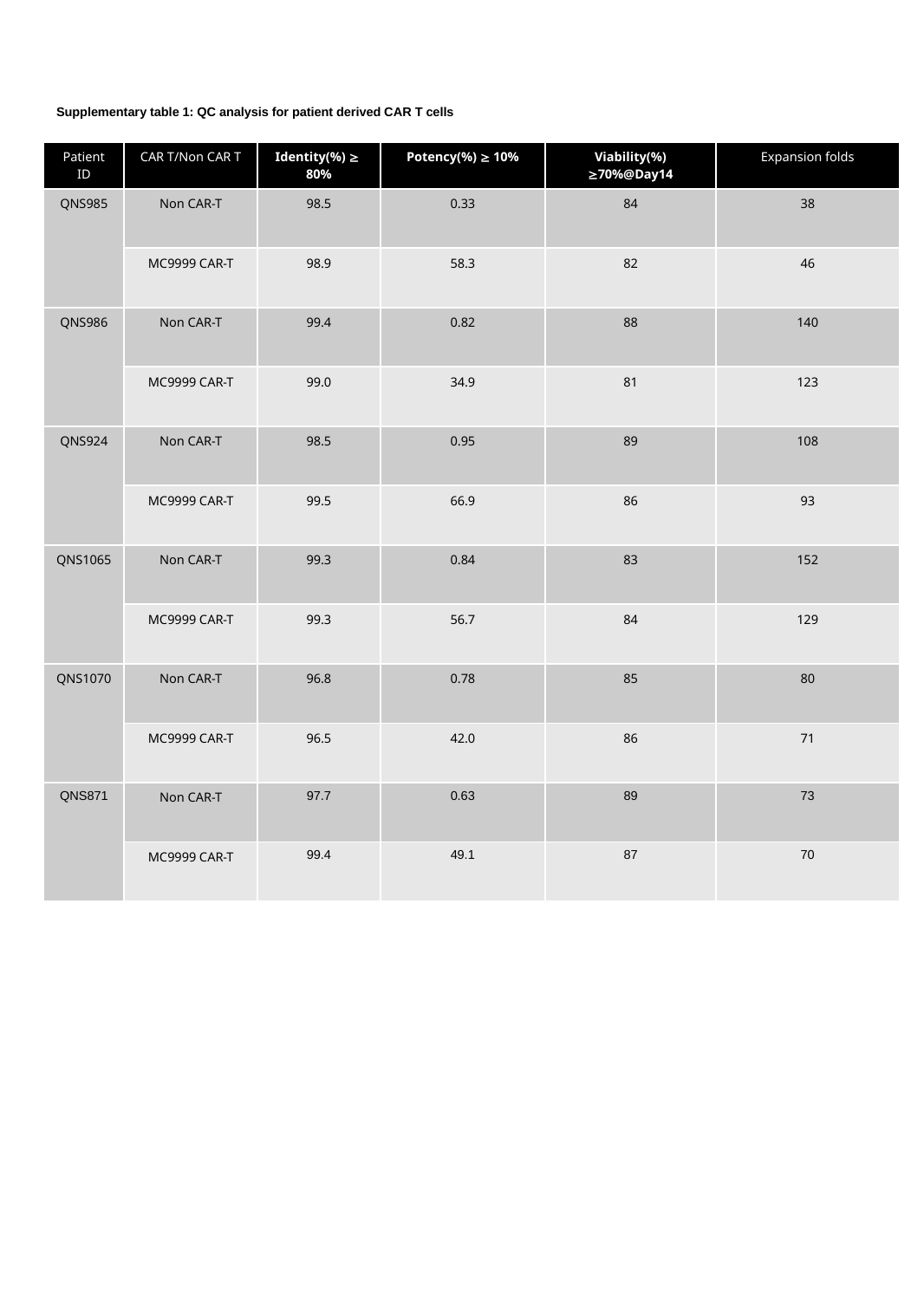

Supplementary table 1: QC analysis for patient derived CAR T cells
| Patient ID | CAR T/Non CAR T | Identity(%) ≥ 80% | Potency(%) ≥ 10% | Viability(%) ≥70%@Day14 | Expansion folds |
| --- | --- | --- | --- | --- | --- |
| QNS985 | Non CAR-T | 98.5 | 0.33 | 84 | 38 |
| | MC9999 CAR-T | 98.9 | 58.3 | 82 | 46 |
| QNS986 | Non CAR-T | 99.4 | 0.82 | 88 | 140 |
| | MC9999 CAR-T | 99.0 | 34.9 | 81 | 123 |
| QNS924 | Non CAR-T | 98.5 | 0.95 | 89 | 108 |
| | MC9999 CAR-T | 99.5 | 66.9 | 86 | 93 |
| QNS1065 | Non CAR-T | 99.3 | 0.84 | 83 | 152 |
| | MC9999 CAR-T | 99.3 | 56.7 | 84 | 129 |
| QNS1070 | Non CAR-T | 96.8 | 0.78 | 85 | 80 |
| | MC9999 CAR-T | 96.5 | 42.0 | 86 | 71 |
| QNS871 | Non CAR-T | 97.7 | 0.63 | 89 | 73 |
| | MC9999 CAR-T | 99.4 | 49.1 | 87 | 70 |
