## Supplementary table 2: Patient clinical and demographic description for "Single CAR-Dual target: Intracranial Delivery of Anti-PD-L1 CAR T Cells Effectively Eradicates Glioma and Immunosuppressive Cells in the Tumor Microenvironment"

### Slide 1
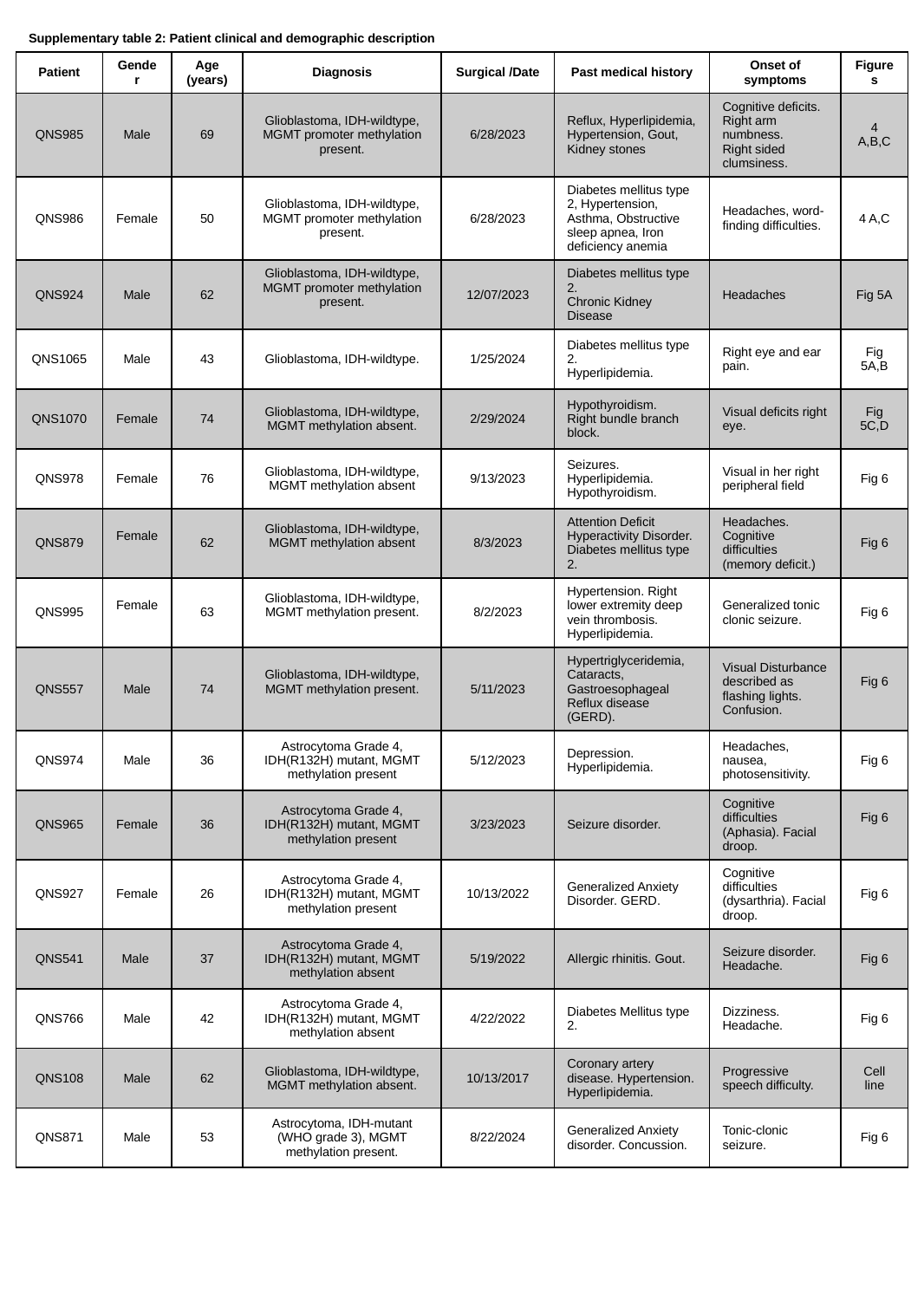

Supplementary table 2: Patient clinical and demographic description
| Patient​ | Gender​ | Age (years)​ | Diagnosis​ | Surgical /Date​ | Past medical history​ | Onset of symptoms​ | Figures​ |
| --- | --- | --- | --- | --- | --- | --- | --- |
| QNS985​ | Male​ | 69​ | Glioblastoma, IDH-wildtype, MGMT promoter methylation present.​ | 6/28/2023​ | Reflux, Hyperlipidemia, Hypertension, Gout, Kidney stones​ | Cognitive deficits.​ Right arm numbness.​ Right sided clumsiness. ​ | 4 A,B,C​ |
| QNS986​ | Female​ | 50​ | Glioblastoma, IDH-wildtype, MGMT promoter methylation present.​ | 6/28/2023​ | Diabetes mellitus type 2, Hypertension, Asthma, Obstructive sleep apnea, Iron deficiency anemia ​ | Headaches, word-finding difficulties.​ | 4 A,C​ |
| QNS924​ | Male​ | 62​ | Glioblastoma, IDH-wildtype, MGMT promoter methylation present.​ ​ | 12/07/2023​ | Diabetes mellitus type 2.​ Chronic Kidney Disease​ | Headaches​ | Fig 5A​ |
| QNS1065​ | Male​ | 43​ | Glioblastoma, IDH-wildtype.​ | 1/25/2024​ | Diabetes mellitus type 2.​ Hyperlipidemia.​ | Right eye and ear pain.​ | Fig 5A,B​ |
| QNS1070​ | Female​ | 74​ | Glioblastoma, IDH-wildtype, MGMT methylation absent.​ | 2/29/2024​ | Hypothyroidism.​ Right bundle branch block.​ | Visual deficits right eye.​ | Fig 5C,D​ |
| QNS978​ | Female​ | 76​ | Glioblastoma, IDH-wildtype, MGMT methylation absent​ | 9/13/2023​ | Seizures. Hyperlipidemia. ​ Hypothyroidism. ​ | Visual in her right peripheral field​ | Fig 6​ |
| QNS879​ | Female​ ​ | 62​ | Glioblastoma, IDH-wildtype, MGMT methylation absent​ ​ | 8/3/2023​ | Attention Deficit Hyperactivity Disorder. Diabetes mellitus type 2.​ | Headaches. Cognitive difficulties (memory deficit.)​ | Fig 6​ |
| QNS995​ | Female​ ​ | 63​ | Glioblastoma, IDH-wildtype, MGMT methylation present.​ ​ | 8/2/2023​ | Hypertension. Right lower extremity deep vein thrombosis. Hyperlipidemia.​ | Generalized tonic clonic seizure. ​ | Fig 6​ |
| QNS557​ | Male​ | 74​ | Glioblastoma, IDH-wildtype, MGMT methylation present.​ ​ | 5/11/2023​ | Hypertriglyceridemia, Cataracts, Gastroesophageal Reflux disease (GERD).​ | Visual Disturbance described as flashing lights. Confusion.​ | Fig 6​ ​ |
| QNS974​ | Male​ | 36​ | Astrocytoma Grade 4, IDH(R132H) mutant, MGMT methylation present​ | 5/12/2023​ | Depression. Hyperlipidemia.​ | Headaches, nausea, photosensitivity.​ | Fig 6​ |
| QNS965​ | Female​ | 36​ | Astrocytoma Grade 4, IDH(R132H) mutant, MGMT methylation present​ | 3/23/2023​ | Seizure disorder.​ | Cognitive difficulties (Aphasia). Facial droop.​ | Fig 6​ ​ |
| QNS927​ | Female​ | 26​ | Astrocytoma Grade 4, IDH(R132H) mutant, MGMT methylation present​ | 10/13/2022​ | Generalized Anxiety Disorder. GERD. ​ | Cognitive difficulties (dysarthria). Facial droop.​ | Fig 6​ |
| QNS541​ | Male ​ | 37​ | Astrocytoma Grade 4, IDH(R132H) mutant, MGMT methylation absent​ | 5/19/2022​ | Allergic rhinitis. Gout.​ | Seizure disorder. Headache.​ | Fig 6​ |
| QNS766​ | Male​ | 42​ | Astrocytoma Grade 4, IDH(R132H) mutant, MGMT methylation absent​ | 4/22/2022​ | Diabetes Mellitus type 2.​ | Dizziness. Headache.​ | Fig 6​ |
| QNS108​ | Male​ | 62​ | Glioblastoma, IDH-wildtype, MGMT methylation absent.​ | 10/13/2017​ | Coronary artery disease. Hypertension. Hyperlipidemia.​ | Progressive speech difficulty.​ | Cell line​ |
| QNS871 | Male | 53 | Astrocytoma, IDH-mutant (WHO grade 3), MGMT methylation present. | 8/22/2024 | Generalized Anxiety disorder. Concussion. | Tonic-clonic seizure. | Fig 6​ |
