## Supplementary figure 2. H&E staining of orthotopic mice models at experimental endpoint. A. Representative section of mice tumors from LN229 cell inje for "Single CAR-Dual target: Intracranial Delivery of Anti-PD-L1 CAR T Cells Effectively Eradicates Glioma and Immunosuppressive Cells in the Tumor Microenvironment"

### Slide 1
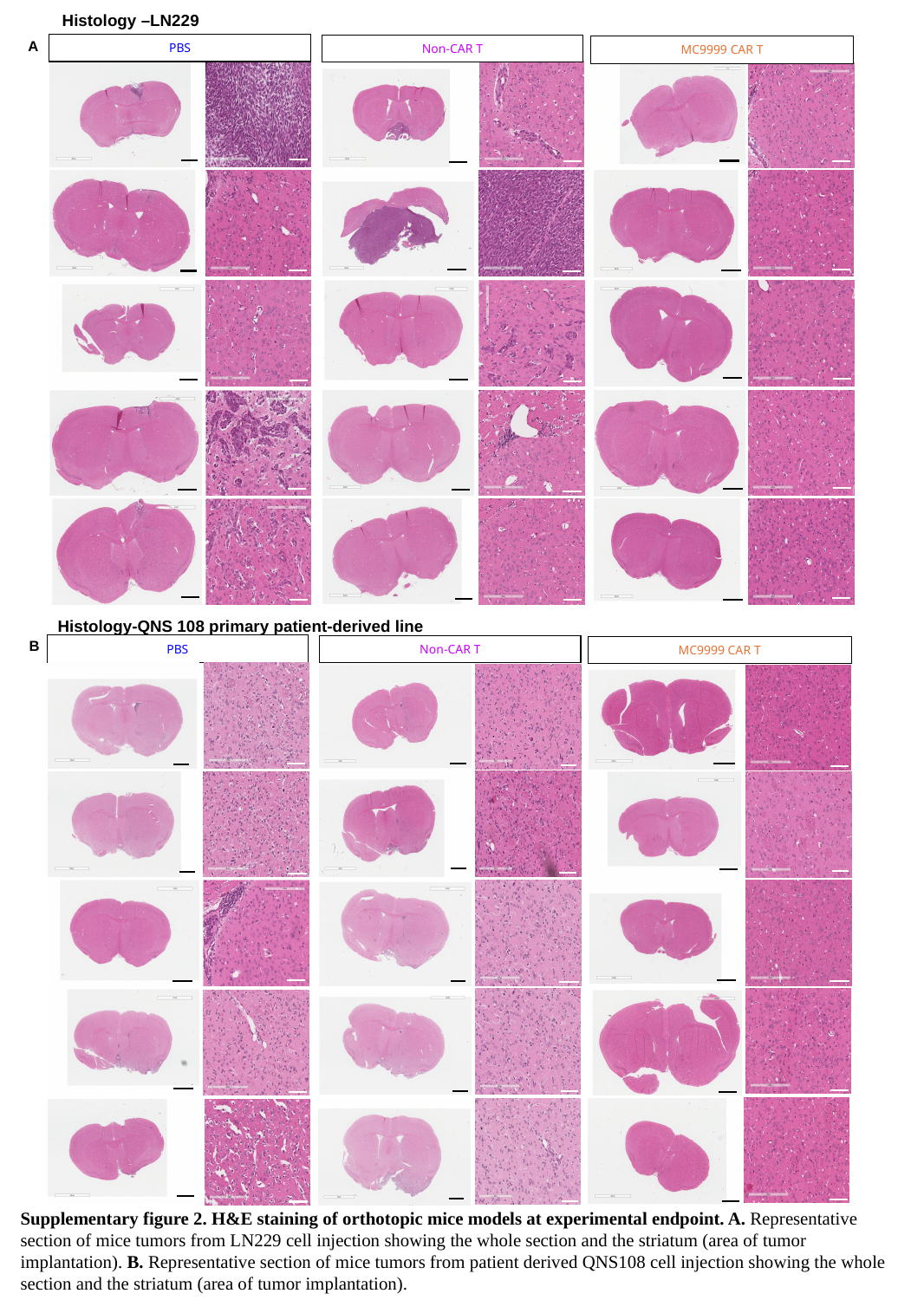

Histology –LN229
PBS
Non-CAR T
MC9999 CAR T
A
Histology-QNS 108 primary patient-derived line
PBS
Non-CAR T
B
MC9999 CAR T
Supplementary figure 2. H&E staining of orthotopic mice models at experimental endpoint. A. Representative section of mice tumors from LN229 cell injection showing the whole section and the striatum (area of tumor implantation). B. Representative section of mice tumors from patient derived QNS108 cell injection showing the whole section and the striatum (area of tumor implantation).
