## Supplementary Figure 5. Timelapse video of CAR-T cell -mediated killing of autologous GBM cells (QNS986) A. incubated with MC9999 CAR T cells; B. Incu for "Single CAR-Dual target: Intracranial Delivery of Anti-PD-L1 CAR T Cells Effectively Eradicates Glioma and Immunosuppressive Cells in the Tumor Microenvironment"

### Slide 1
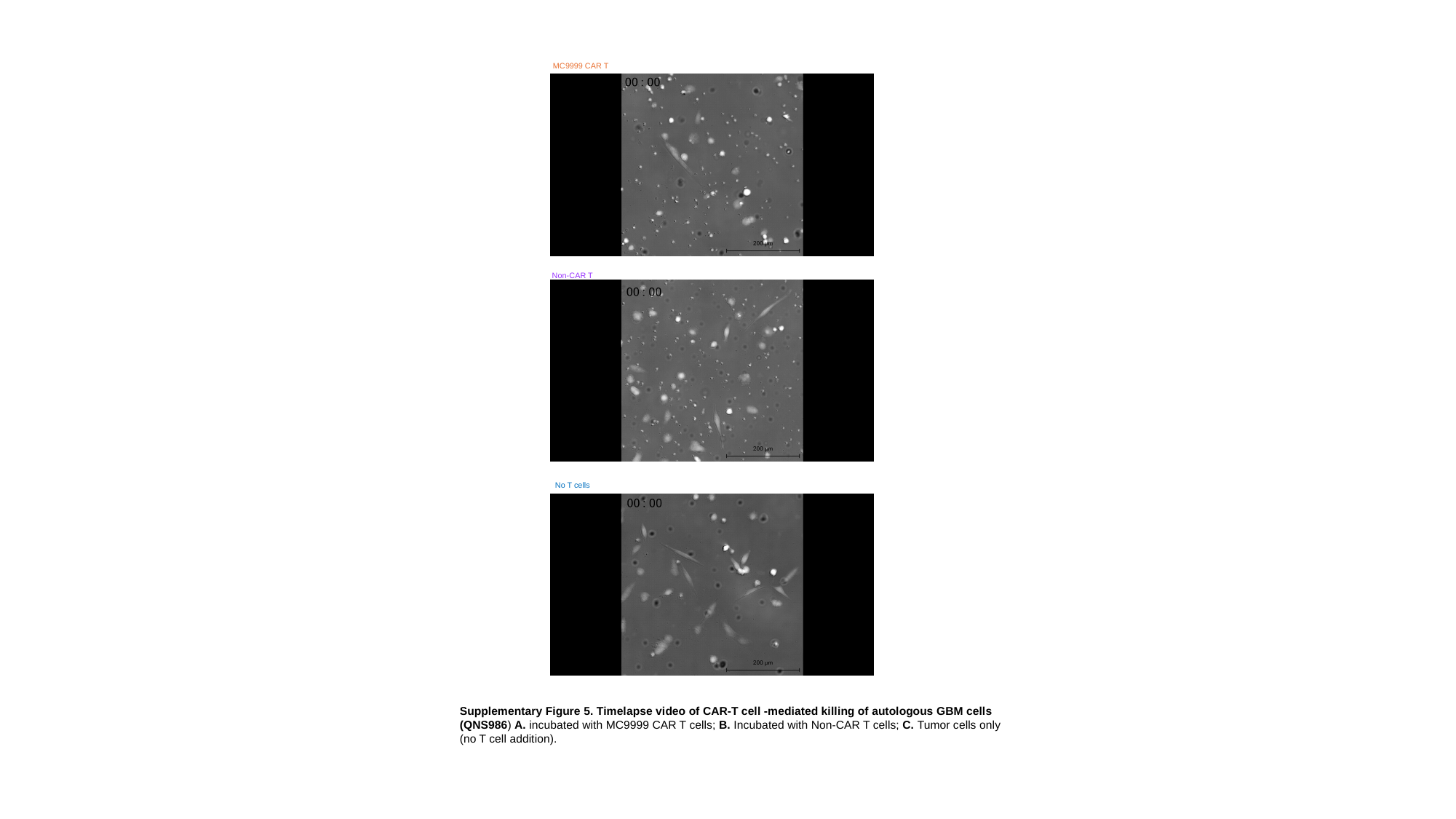

MC9999 CAR T
Non-CAR T
No T cells
Supplementary Figure 5. Timelapse video of CAR-T cell -mediated killing of autologous GBM cells (QNS986) A. incubated with MC9999 CAR T cells; B. Incubated with Non-CAR T cells; C. Tumor cells only (no T cell addition).
